# The glutathione S-transferase Gstt1 is a robust driver of survival and dissemination in metastases

**DOI:** 10.1101/2022.09.10.507413

**Authors:** Christina M. Ferrer, Ruben Boon, Hyo Min Cho, Tiziano Bernasocchi, Lai Ping Wong, Murat Cetinbas, Elizabeth R. Haggerty, Irene Mitsiades, Gregory R. Wojtkiewicz, Daniel E. McLoughlin, Sita Kugel, Esther Rheinbay, Ruslan Sadreyev, Dejan Juric, Raul Mostoslavsky

## Abstract

Identifying adaptive mechanisms of metastatic cancer cells remains an elusive question in the treatment of metastatic disease, particularly in pancreatic cancer (PDA), where the majority of patients present with metastatic lesions at the time of diagnosis. A loss-of-function shRNA targeted screen in metastatic-derived cells identified *Gstt1*, a member of the glutathione S-transferase superfamily, as uniquely required for metastasis and dissemination however dispensable for primary tumor growth. *Gstt1* is expressed in early disseminated tumor cells (DTCs), is retained within a subpopulation of slow-cycling cells within established metastases and its inhibition led to a regression of macrometastatic lesions. This distinct Gstt1^high^ population is highly metastatic and retains slow-cycling phenotypes, EMT features, and DTC characteristics compared to the Gstt1^low^ population. Mechanistic studies indicate that in this subset of cells, Gstt1 maintains metastases by binding to and modifying intracellular fibronectin, regulating Fibronectin secretion from cancer cells and deposition into the metastatic microenvironment. We identified Gstt1 as a novel mediator of metastasis, highlighting the importance of metastatic heterogeneity and its influence on the metastatic tumor microenvironment.

## INTRODUCTION

Only 0.01% of cancer cells enter circulation, survive and produce metastasis, however, metastatic disease accounts for 90% of cancer-related deaths (*5,10,48*). In order to establish metastases, cancer cells from the primary tumor navigate numerous hurdles: traversing blood vessels, surviving in the circulation, and colonizing and propagating at foreign habitats from single cells or small groups of cells (*9,10,23*). Thus, distinct molecular programs are required to endow cancer cells with the ability to adapt to the metastatic niche and develop overt metastases. Notably, much of the research into the mechanisms of metastasis has been focused on identifying early drivers within primary tumors. In pancreatic adenocarcinoma (PDA), several studies have shown that metastatic lesions lack additional driver mutations compared to primary tumors (*27, 28,55*), and thus, elucidating the circuitry distinguishing metastases from primary tumors is an urgent question for the field. Although specific epigenetic and stage-dependent adaptations have been identified between metastases and primary tumors (*26,29,36,37,39,41,51*), few studies have addressed the importance of metastatic heterogeneity in pancreatic cancer (*19,27,31,36*).

It is widely understood that metastatic dissemination is an early event in cancer progression (*5,9,17,20,21,23*). This is particularly evident in PDA where the majority of patients present with extra-pancreatic invasion and metastatic disease upon diagnosis of the primary lesion, for which five-year survival rate is only ∼7.1% (*38,42,46*). Thus, interventions aimed at targeting mechanisms of primary tumor dissemination are unlikely to be clinically successful strategies owing to the presence of synchronous or undetectable metastases (*17,20,21,36*). Additionally, the biological heterogeneity observed amongst primary tumors, disseminated disease and overt metastases, as well as our limited understanding of these processes presents a clinical challenge and an urgent question in the field. While some advancements have been made in understanding the mechanisms of metastatic dissemination (*38, 51*), latency (*36*), and outgrowth (*26,27,29,31,41*), these topics remain largely unexplored in PDA.

Utilizing orthotopic models of metastasis, we have performed unbiased RNA-Seq on matched primary and metastatic solid tumors from established mouse models of PDA and breast cancer. Sequencing analysis was followed by a targeted functional shRNA screen to identify genes conferring a 3D growth advantage in metastasis-derived cells. In this study, we provide evidence that *Gstt1*, the gene encoding a glutathione S-transferase with no known roles in cancer, is uniquely expressed in and required for the formation of macrometastases while remaining dispensable for primary tumor growth. Within metastatic lesions, Gstt1 displays a heterogenous expression pattern, where Gstt1^high^ cells represent a highly metastatic, slow-cycling population with EMT features. Importantly, this slow-cycling, Gstt1^high^ subpopulation is not only retained in established lesions, but is also required for metastatic dissemination. Mechanistically, we find that Gstt1 directly glutathione-modifies intracellular Fibronectin, thus enhancing the secretion of Fibronectin from tumor cells and its deposition into the extracellular matrix (ECM), which appears to be critical for maintaining metastatic growth. Overall, we identified Gstt1 as a novel metastatic driver in a subpopulation of metastatic cells and highlight the interplay between heterogeneity within tumor cells as critical in shaping the metastatic microenvironment.

## RESULTS

### RNA-Seq Identifies an Upregulated Gene Signature in Pancreatic and Breast Cancer Metastases

Since most of the previous screening methods have focused on early initiating events in metastasis (*5,14,48*), we sought to explore whether we could identify genes required for ‘metastatic maintenance’ using established mouse models based on transcriptional differences between primary tumors and macrometastases. We isolated tissue from matched primary and metastatic tumors from two genetically defined mouse models of pancreatic ductal adenocarcinoma (PDAC) (*p48-Cre/p53F/+KrasL/+* (*Sirt6 WT* and *KO*) which closely mimic progression of human primary pancreatic cancer and generate spontaneous distant metastases to the liver and lung, with the more aggressive *Sirt6 KO* model readily developing lung metastases (*22,41,42*) (**Figure 1A, left panel**). To identify more general transcriptional adaptations across carcinoma-derived metastases, we included the 4T1 breast cancer (BC) mouse model which develops spontaneous metastases to the lung when orthotopically injected into the mammary fat pad of Balb/c mice (*6*) (**Figure 1A, right panel**). Together, these models provide spontaneous metastases in an immune competent setting while preserving the steps of the metastatic cascade. We performed RNA-Seq on the isolated bulk tissue primary tumors and metastases to generate expression profiles corresponding to PDAC (n=4) and BC (n=4) primary tumors, as well as lung (n=8) and liver metastases (n=4) (**Figure 1A**). Subsequent analysis identified a common differentially upregulated gene signature (n=136) in distant metastases (liver and lung) compared to matched primary tumors (2FC, FDR 0.05) across all tumor types (**Figure 1B, 1C**).

**Figure 1.**
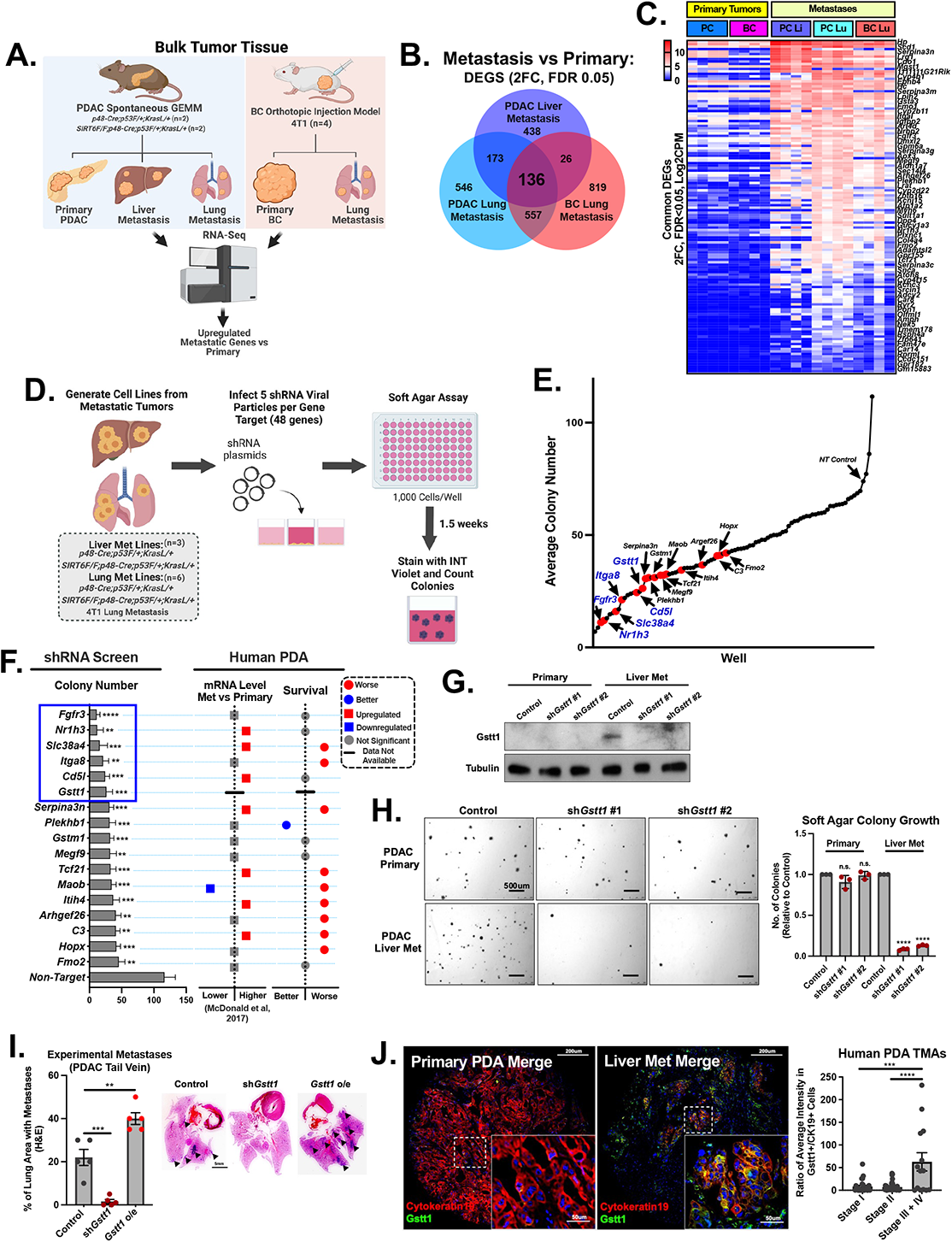
(A) Diagram depicting mouse models used for metastasis experiments. (Left) PDAC transgenic model (*p48-Cre/p53F/+KrasL/+* (*SIRT6* WT and KO)) generates spontaneous locally invasive disease (lymph node) (n=4) and distant metastasis to the liver (n=4) and lung (n=4). (Right) Orthotopic allograft injection of 4T1 mouse breast cancer cell line (BC) results in primary tumor growth (n=4) and spontaneous lung metastasis (n=4). Bulk tissue primary tumors and metastases were subjected to RNA-Seq analysis. (B) Venn Diagram demonstrating common and unique differentially expressed genes between metastatic and primary tumors. N=136 genes are commonly differentially expressed in all metastases vs matched primary tumors (Log2FC, FDR 0.05). (C) Heatmap of 136 commonly differentially expressed genes in all metastases and primary tumors expressed as Log2CPM (Log2FC, FDR 0.05). (D) Schematic of targeted soft agar 96-well shRNA screen in metastasis-derived cells. One individual gene target is represented in each well. (E) shRNA target genes ranked by average colony number per well (lowest to highest) compared to control shRNA. Wells displaying significantly reduced colony growth represented as red dots (Top 17). Top 6 candidate genes labeled in blue. (F) Top 17 shRNA target genes in order of average colony number (Left). *t-test* was used to determine statistical significance between non-targeting control and average colony number for each shRNA (p**<0.01; p<***<0.05; p****<0.001). Top 17 shRNA target genes were evaluated for differential expression in human PDA metastases compared to matched primary tumors (Upregulated in mets, red; Downregulated in mets, blue; Not Significant in gray) (McDonald et al, 2017, n=16 primary, liver met and lung metastatic samples) (Middle). Top 17 genes were evaluated using human PDA TCGA datasets to determine prognostic consequences of high gene expression on progression free survival (worse prognosis, red; better prognosis, blue; not significant in gray) (Right). (G) Western blot depicting PDAC-derived primary and metastatic cell lines stably expressing control or two independent *Gstt1* shRNAs. (H) Soft agar assay growth for *Gstt1* in matched primary and metastatic cell lines. Represented as number of colonies per well relative to control (p****<0.0001) (Right) Representative bright-field image (2.5X) (Left). The experiment was performed in triplicate with three replicates each. Data are mean s.d. Student’s *t-test* was used to determine statistical significance between groups (p****<0.0001). (I) SCID mice were injected via tail vein with liver metastatic (PDAC) cell lines stably expressing Control, sh*Gstt1* or doxycycline inducible *Gstt1* cDNA and sacrificed after 4 weeks. Left panel. Lung metastatic burden was evaluated using H&E and represented as total lung tissue area versus the metastatic tissue area. Data are represented as mean s.e.m. *t-test* was used to determine statistical significance between groups (p**<0.01, p***<0.001). Right panel. Representative H&E images of lung quantification from left panel. (J) Immunofluorescence staining of Cytokeratin 19 and Gstt1 in human PDA Tissue Microarray. Representative image of primary tumor core and liver met core. Quantification of all cores (Stage I and II n= 68 cores; Stage III and IV n=14). *t-test* was used to determine statistical significance between groups (p***<0.001; p****<0.0001).

### Targeted Functional shRNA Screen to Identify Genes Required for Metastatic-Cell Growth

In order to functionally test whether this upregulated ‘metastatic signature’ was required for the growth and survival of established metastatic cells, we developed a targeted, functional 96-well shRNA screening method based on anchorage-independent growth (**Figure 1D**). Using our mouse models of metastasis, we generated metastases-derived cell lines from spontaneous tumors originating in the liver (PDAC, n=3) and lung (PDAC, n=3; BC, n=3). Employing individual shRNAs targeting each gene (top 94 differentially expressed genes x 5 pooled shRNAs per target), metastatic-derived cell lines were infected with 5 pooled shRNA viral particles in a 96-well format to establish stably expressing cells. Metastatic cells were then subjected to a soft agar colony formation assay in a 96-well format (n=9 total cell lines/gene target). After two weeks, soft agar colonies were stained with INT violet, counted and photographed (**Figure 1D**). Wells were ranked by colony number from lowest to highest (**Figure 1E**). Wells displaying a statistically significant reduction in soft agar colony growth (indicated as red dots) were identified as genes required for anchorage-independent growth of metastatic cells. Using this approach, we identified 17 genes that impaired 3D growth upon depletion (**Figure 1E**). To validate differential expression of the 17 candidate genes and account for surrounding stromal or microenvironment contamination, we performed a parallel analysis on tumor cell-intrinsic expression in lineage-traced YFP/GFP+ primary and metastatic cells. To isolate primary and metastatic tumor cells, we utilized our genetic PDAC mouse models (*p48-Cre/p53F/+KrasL/+/ROSA26-LSL-YFP* (*Sirt6 WT* and *KO*) (n=2) and 4T1 cells tagged with GFP injected orthotopically into the mammary fat pad (n=2). Mice were euthanized when both groups showed visible clinical signs of metastasis. Primary tumor and metastatic tissues were digested and YFP and GFP-lineage-traced cells were isolated using fluorescence activated cell sorting (FACS) followed by RNA sequencing on the purified cell populations to generate gene expression profiles devoid of stromal contamination (**Figure S1A**). RNA-Seq performed on GFP+ lineage-traced metastases revealed that 3 out of the 17 genes identified in our screen (*Nr1h3*, *Gstm1*, *Itih4*) displayed differential expression in metastatic tumor cells compared to primary tumor cells with others (*Cd5l*, *Gstt1*, *C3*, *Hopx*) showing an expression trend similar to results observed in our bulk tissue analysis (**Figure S1B**). To further validate metastasis-specific genes emerging from our screen, we analyzed differential expression of our top 6 candidate genes (**Figure 1E, 1F, blue**) in our cell line screening model, which demonstrated enrichment in *Fgfr3*, *Nr1h3*, *Slc38a4* and *Gstt1* in metastatic cell lines compared to primary tumor-derived lines (**Figure S1C**). We then validated our top 6 candidates by examining the effect of individual shRNAs on soft agar growth (**Figure S1D, S1E**). Inhibiting *Nr1h3*, *Slc38a4, Cd5l* and *Gstt1* verified our original screen results, with two independent shRNAs significantly affecting metastatic cell growth in 3D (**Figure S1E, S1F**).

To continue to narrow down metastasis-specific candidate genes emerging from our screen and assess their relevance in human disease, we examined their expression in metastatic vs primary PDA (*24, 29*) and determined whether high expression was associated with poor prognosis using publicly available TCGA datasets. 5 out of 17 genes identified in our screen (*Slc38a4*, *Serpina3n*, *Tcf21*, *Itih4*, *C3*) demonstrated both criteria: differential enrichment in PDA metastases and worse prognosis (RFS) in human PDA (**Figure 1F**) (*24, 29*). Notably, several of these genes have been previously shown to be well-defined mediators of metastatic adaptation in other tumor types (*5,8,53*), suggesting some mutually conserved mechanisms for metastasis across primary tumors. Interestingly, from our top 6 validated candidate genes, *Slc38a4* demonstrated enrichment in these criteria, however, *Gstt1* was largely absent from pancreatic cancer datasets (**Figure 1F**, **Figure S1G**), presenting an intriguing question to explore as to the molecular roles for *Gstt1* in cancer, and, in the context of metastatic disease.

Further validation of our screening strategy confirmed *Gstt1* as being required for soft agar colony growth (**Figure 1H, S1E**), dispensable for 2D colony growth (**Figure S1J**) and uniquely required for tumor sphere forming potential (**Figure S1H, S1I**) of metastatic-derived cells *in vitro*. Strikingly, cells derived from primary PDAC tumors do not show dependence on *Gstt1* for anchorage-independent growth, consistent with to the low expression levels observed in primary tumors (**Figure 1G, 1H**).

### The Glutathione Conjugating Enzyme *Gstt1* is Required for Metastatic Growth *in vivo*

*Gstt1* encodes the glutathione S-transferase (GST) theta 1 enzyme, a member of a superfamily of proteins that catalyze the conjugation of reduced glutathione (GSH) to a variety of electrophilic and hydrophobic compounds, including carcinogens, chemotherapeutic drugs, and many oxidative metabolic byproducts (*50, 52*). Consequently, GSTs play a critical role as cellular detoxifying enzymes by predisposing compounds to further modification and clearance, specifically in the liver. Notably, the *Gstt1* gene is haplotype specific and absent from ∼30% of the population (*3,50*,*52*), suggesting redundant functions in normal cells. Although various groups have reported disparate roles for the association between *Gstt1* null polymorphism as a risk factor for the development of cancer, molecular roles for Gstt1 in cancer, and specifically, in the context of metastatic disease, remain unknown (*3*).

To validate dependences *in vivo*, we sought to determine whether *Gstt1* is both required and whether it can enhance the formation of metastasis. Using our experimental PDAC model, lung-metastatic derived cell lines stably expressing either non-targeting control, *Gstt1* shRNA or a *Gstt1* doxycycline-inducible overexpression construct (**Figure S1K**) were injected via tail vein and euthanized 4 weeks post-injection. Strikingly, we observed a significant reduction in metastatic lesions in our *Gstt1* shRNA expressing cells (**Figure 1I**). Conversely, overexpressing *Gstt1* in metastatic cells lead to increased metastatic burden in tail vein injected mice (**Figure 1I**), pointing to *Gstt1* as a novel driver of metastatic potential in PDAC.

To validate metastatic dependencies in both tumor models, we utilized both experimental and spontaneous models of metastasis using cell lines derived from our 4T1 tumors. To test *Gstt1* as being required for experimental metastasis, lung-metastatic derived lines from our BC model were injected into the venus sinus (retroorbital) to measure the formation of lung metastases (**Figure S1L**). Notably, inhibition of Gstt1 almost completely abolished the formation of metastatic lesions (**Figure S1M**). As a model of spontaneous metastasis, primary-derived cell lines from our 4T1 BC model stably expressing sh*Gstt1* (**Figure S1L**) were injected orthotopically into the mammary fat pad (1 x 10^5^ cells), and mice were monitored for both primary tumor size and the formation of end-stage metastases (mice sacrificed at 4 weeks post-injection due to primary tumor burden). Remarkably, while the growth of primary tumors was unaffected (**Figure S1N**), knockdown of *Gstt1* resulted in a significantly reduced number of spontaneous metastatic lesions (**Figure S1O, S1P**). Together with our *in vitro* data, these results point support the robustness of our screening strategy together pointing to a unique requirement for *Gstt1* specifically in the context of metastatic disease.

### Patient-derived Pancreatic Metastases Display Increased Expression and Dependence on Gstt1 for Metastatic Growth *in vitro*

As previously mentioned, PDAC GEM models closely recapitulate the full scope of human pancreatic cancer, including metastatic disease (*1,22,31*). We next examined the expression of Gstt1 across metastatic tissue types in our mouse models of PDAC. Confirming our RNA-Seq results and expression in metastatic cell lines, immunofluorescence imaging of CK19-positive PDAC primary tumor cells and matched metastases demonstrated that Gstt1 is expressed in a subset of metastatic lesions and widely absent from primary tumors (**Figure S2A, S2B**). As previously noted, the dearth of publicly available datasets for examining *Gstt1* presented a specific hurdle to characterizing this gene in pancreatic cancer. To overcome this, we collected and analyzed a panel of matched human primary and metastatic PDA lesions collected via rapid autopsy. We began our analysis by immunostaining formalin-fixed patient tissue samples derived from primary tumors, liver, and lung metastases (multiple lesions from 3 patients). For each patient, tissue samples were taken from different regions of the primary tumor (if present at post-mortem) as well as matched metastatic lesions. Our analysis revealed upregulation of Gstt1 across metastatic samples relative to primary tumors by both immunofluorescence staining of formalin-fixed tissue sections (**Figure S2C**) and bulk tissue western blot (**Figure S2D**). To add to the robustness of the analysis, we obtained a pancreatic cancer tissue microarray (TMA) panel containing both primary PDA cores as well as metastatic cores and performed immunofluorescence for Gstt1. We observed a striking enrichment for Gstt1 in CK19-positive metastatic cores (Stage III/IV, n=14) compared to primary tumor-derived cores (Stage I/II, n=68) (**Figure 1J**). To further validate the functional significance of Gstt1 in human PDA, we also silenced *Gstt1* in two metastatic-derived human pancreatic cancer cell lines, KP3 and CFPAC1 (**Figure S2E**). Inhibiting *Gstt1* resulted in a significant reduction in soft agar colony formation in liver metastatic cell lines (**Figure S2F**). These observations provide strong support for our screening strategy in the mouse and identified Gstt1 as being differentially upregulated in metastatic PDA, thus potentially contributing to survival outcomes in these patients.

### Gstt1 is Required for the Maintenance and Dissemination of Metastatic Cells in PDAC

To assess the role of Gstt1 in metastatic maintenance, we employed an inducible Cas9 system to target *Gstt1* in mice with established metastases. Lung metastatic-PDAC derived iCas9 control and sg*Gstt1* expressing cells (**Figure 2A)** were injected retro-orbitally to achieve lung metastases. Mice were monitored via bioluminescence imaging (BLI) for evidence of metastases. Upon evidence of metastatic signal, iCas9 and iCas9 + sg*Gstt1* injected mice were treated with doxycycline in the drinking water, followed by continuous monitoring using bioluminescence. Strikingly, bioluminescence imaging of lung metastases revealed a decrease in signal in the iCas9 + sg*Gstt1* mice as early as 7 days post-doxycycline, while iCas9 control expressing mice continued to progress and eventually succumbed to metastatic disease (**Figure 2B, 2C**), demonstrating that *Gstt1* is not only required for the formation of metastasis but also critical in the maintenance of established tumors.

**Figure 2.**
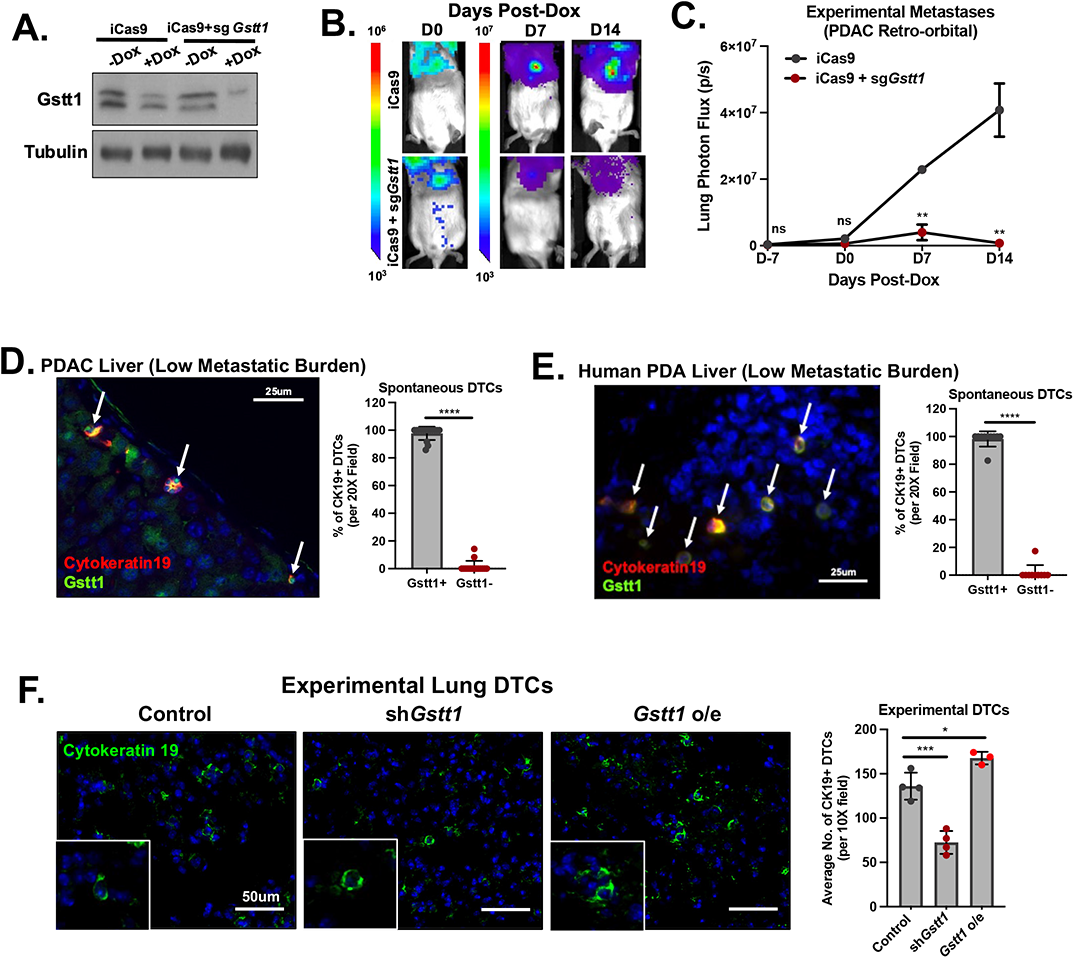
(A) SCID mice were injected retro-orbitally with PDAC lung metastatic-derived cells expressing iCas9 Control and iCas9 + sg*Gstt1*. Western blot demonstrating dox-inducible KO of *Gstt1*. (B) Representative bioluminescent images of pre-Dox (D0) and post-Dox (D7, D14) metastatic burden. (C) Quantified lung photon flux pre-Dox (D7, D0) and post-Dox (D7, D14). Data are represented as mean s.e.m. *t-test* was used to determine statistical significance between groups (p**<0.01). (D) Immunofluorescence staining of Gstt1 and Cytokeratin 19 (pancreatic cell marker) in PDAC-derived liver disseminated tumor cells (DTCs) from low tumor burden mice. Quantification represents the average of 10 fields from a minimum of n=3 mice. Data are represented as mean s.d. *t-test* was used to determine statistical significance between groups (p****<0.0001). (E) Immunofluorescence staining of Gstt1 and pan-Cytokeratin (tumor cell marker) in PDA-derived liver disseminated tumor cells (DTCs). Quantification represents the average of 10 fields for PDA patient #4. Data are represented as mean s.d. *t-test* was used to determine statistical significance between groups (p****<0.0001). (F) Immunofluorescence staining of experimental disseminated tumor cells (DTCs) using Cytokeratin 19 2 weeks post-injection. Arrows indicate DTCs and inset panel demonstrates a magnified image of DTCs. Quantification represents the average of 10 fields from a minimum of n=3 mice. Data are represented as mean s.d. *t-test* was used to determine statistical significance between groups (p*<0.05; p***<0.005).

Similar metastasis-specific dependencies were observed in 4T1 cells expressing an inducible Cas9 cassette (iCas9), where doxycycline was given prior to orthotopic cell injection thereby inhibiting *Gstt1* beginning at tumor cell implantation (**Figure S3A**), followed by removal of the primary tumor and monitoring of metastasis using BLI (*11*). Inducible inhibition of *Gstt1* from D0 post-implantation did not affect the growth of the orthotopic primary tumor (**Figure S3B**), while resulting in mice devoid of lung metastases, compared to iCas9 control animals (**Figure S3C-S3E**), pointing to a potential role for Gstt1 in the metastatic process at stages after dissemination from the primary tumor.

To further test the specificity of Gstt1 in mediating metastasis *in vivo*, we performed rescue experiments in control and sh*Gstt1* expressing cells by overexpressing a doxycycline-inducible *Gstt1* cDNA in primary-tumor derived cells from the 4T1 BC mouse model (**Figure S3F)**. Specifically, mice were treated with doxycycline in the drinking water 7 days prior to orthotopic cell injection into the mammary fat pad (1 x 10^5^ cells) to induce *Gstt1* expression during tumor cell implantation, followed by BLI to monitor formation of metastases upon surgical removal of the primary tumors. Re-expression of *Gstt1* in the context of the sh*Gstt1* background did not affect the formation of primary tumors (**Figure S3G**), further supporting metastasis-specific dependencies. More importantly, reconstitution of *Gstt1* not only rescued the absence of lung metastases upon deletion of *Gstt1* (**Figure S3H-J**), but strikingly, it drove metastases in ectopic regions throughout the mouse (**Figures S3H, S3K, S3L**) resulting in the formation of bone, diaphragm and liver tumors. Together with results observed in our PDAC experimental metastasis model, these results suggest that Gstt1 may play a crucial role in tumor cell dissemination. Although we observed clear phenotypes for *Gstt1* in both pancreatic and breast cancer metastases, in the present study we will focus on pancreatic cancer for further characterization.

### Gstt1 is Expressed in DTCs and is Required for Early Tumor Cell Dissemination

Our results demonstrating the formation of ectopic metastases points to *Gstt1* as a mediator of tumor cell dissemination. Disseminated tumor cells, or DTCs, are characterized as a group of 1-5 cells that have escaped the primary tumor and arrived at a distant organ (*9,16,17,20,21,23,36*). To assess whether *Gstt1* may be important in early dissemination as well, we first assessed Gstt1 expression in DTCs, by examining low burden tissues derived from PDAC GEMM models with primary tumors but no visible macrometastases. Strikingly, our analysis revealed that almost 100% of single DTCs and groups of 1-5 DTCs stained for both CK19 and Gstt1. This was observed in both low metastatic burden tissues from PDAC mice (**Figure 2D**) and in the liver tissue of a PDA-derived rapid autopsy patient presenting with a primary tumor and low metastatic burden (**Figure 2E**).

To determine whether Gstt1 is required for tumor cell dissemination, we inhibited *Gstt1* in our primary-derived PDAC cell lines and performed tail vein injections to derive experimental lung DTCs. To stain for DTCs, we sacrificed the mice at early timepoints (2 weeks post-tail vein injection). Depleting *Gstt1* resulted in significantly fewer disseminated single cells and clusters (**Figure 2F**), and overexpressing *Gstt1* enhanced the number of DTCs (**Figure 2F**). To determine the effect on spontaneous tumor cell dissemination, we inhibited *Gstt1* in our primary-derived PDAC cell lines and performed orthotopic injections into the pancreas, a model that requires euthanasia due to primary tumor burden. As expected, mice injected with primary tumor cells displayed no visible evidence of metastatic burden. Liver and lung tissues were then stained with CK19 to detect disseminated tumor cells. We observed that control treated cells readily disseminated and thus we were able to detect both single and groups of DTCs (**Figure S4C**), which also stained for Gstt1 (**Figure S4B**). However, depleting *Gstt1* resulted in significantly fewer disseminated single cells and clusters (**Figure S4C, S4D**), without affecting primary tumor growth (**Figure S4A**). Together, these results indicate the requirement for *Gstt1* following escape from the primary tumor, allowing for dissemination and survival of DTCs in the metastatic niche.

### Gstt1^high^ DTCs and Established Metastases Represent a Slow-Cycling Subpopulation within Metastases

A growing body of evidence suggests that DTCs undergo an extended period of quiescence in the stroma of the target organ, where they are subjected to a selective process, where rare cells may ‘reawaken’, leading to overt metastases (*15,16,17*). In order to assess the cycling status of these Gstt1-positive cells, we stained for PCNA, a proliferation marker. We found that most of our Gstt1-positive DTCs did not stain for PCNA in both our spontaneous GEMM (**Figure 3A, 3B**) and experimental metastasis models (**Figure S5A, S5B**). Similarly, the DTCs we identified in the liver tissue of our PDA-derived rapid autopsy patient also stained negative for Ki67, another marker of proliferation (**Figure 3C, 3D**).

**Figure 3.**
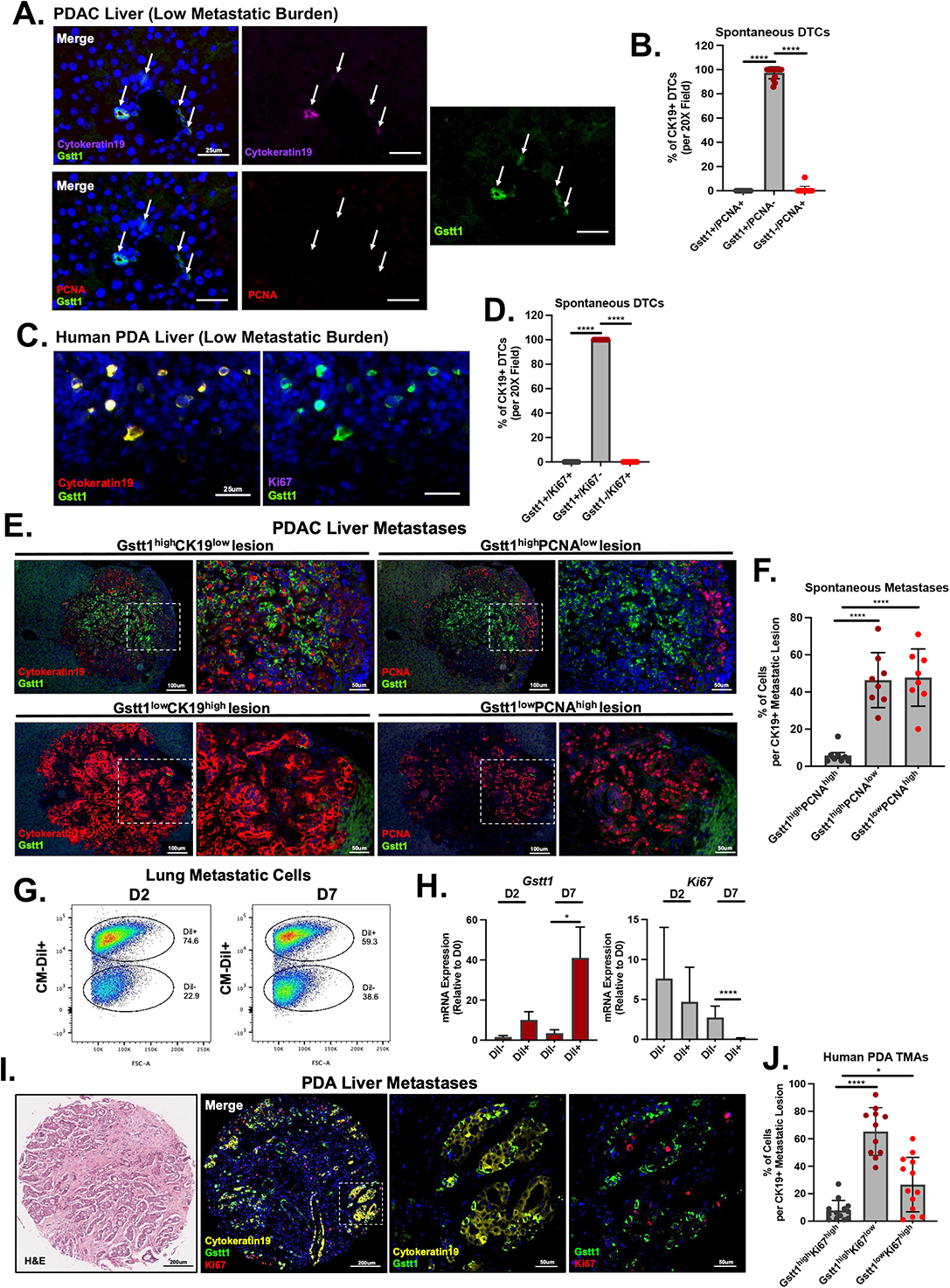
(A) Immunofluorescence staining of Gstt1, Cytokeratin 19 (pancreatic cell marker) and PCNA in PDAC-derived liver disseminated tumor cells (DTCs) from low tumor burden mice. (B) Quantification of (A) represents the average of 10 fields from a minimum of n=3 mice. Data are represented as mean s.d. *t-test* was used to determine statistical significance between groups (p****<0.0001). (C) Immunofluorescence staining of Gstt1, Ki67 and Cytokeratin 19 (pancreatic cell marker) in PDA-derived liver disseminated tumor cells (DTCs). (D) Quantification of (C) represents the average of 10 fields for PDA patient #4. Data are represented as mean s.d. *t-test* was used to determine statistical significance between groups (p****<0.0001). (E) Immunofluorescence staining of Gstt1 and Cytokeratin 19 (pancreatic cell marker) in matched PDAC-derived liver metastatic tissue (left panel). Immunofluorescence staining of Gstt1 and PCNA in PDAC-derived liver metastases (right panel). (F) Quantification of cell populations represents the average of 5 fields per n=8 independent metastatic lesions. Data are represented as mean s.d. *t-test* was used to determine statistical significance between groups (p****<0.0001). (G) FACS analysis of lung metastatic cell populations 2 and 7 days post-incubation with membrane dye CM-DiI. (H) RT-PCR expression for *Gstt1* and *Ki67* in CM-Dil- and CM-Dil+ sorted populations from (G). Data are represented as mean s.e.m. *t-test* was used to determine statistical significance between groups (p*<0.05; p****<0.0001). (I) H&E image (left) and immunofluorescence staining of Cytokeratin 19, Ki67 and Gstt1 in liver metastatic core from human PDA Tissue Microarray (right). (J) Quantification includes omental, abdominal and liver metastatic cores (Stage IV n=10). Data are represented as mean s.d. *t-test* was used to determine statistical significance between groups (p*<0.05; p****<0.0001).

Next, we analyzed our high metastatic burden tissues for Gstt1 and proliferation markers. Intriguingly, immunofluorescence imaging of CK19-positive PDAC primary tumors and matched metastases demonstrated that Gstt1 is not only heterogeneously expressed within individual metastatic lesions but also demonstrates heterogeneity across metastatic lesions (**Figure 3E, left panels, Figure 3F**). Interestingly, this Gstt1^high^ subpopulation in metastatic tumors was associated with a CK19^low^ phenotype, with high burden tissues containing both Gstt1^high^/CK19^low^ lesions as well as Gstt1^low^/CK19^high^ lesions (**Figure 3E, left panels**), a phenomenon previously observed in non-proliferating PDAC DTCs (*36*). Furthermore, staining for proliferation markers revealed that a large majority of Gstt1 positive cells were PCNA-negative (∼95%), suggesting a slow-cycling state in Gstt1^high^ metastatic cells, coexisting with Gstt1^low^/PCNA^high^ subpopulations (**Figure 3E, right panels, Figure 3F**). To further quantify the cycling profile of *Gstt1* within our experimental models, we labeled our PDAC-derived lung metastatic cell lines with a widely used membrane dye, CM-DiI (*25, 33*). *In vitro* cultures of labeled cells demonstrated a dilution of the membrane dye over time, with the accumulation of two distinct cycling populations (**Figure 3G, Figure S5C**). Next, CM-Dil+ and CM-DiI-negative populations were isolated using fluorescence-activated cell sorting (FACS) and analyzed for *Gstt1* and *Ki67* mRNA expression. RT-PCR results demonstrate that CM-DiI+ label-retaining cells at day 7 post-labeling are highly enriched for *Gstt1* expression compared to CM-DiI-cells (**Figure 3H**), and as expected, they exhibit low levels of *Ki67* expression (**Figure 3H**). All together, these results indicate that Gstt1 marks a metastasis-specific, slow-cycling population.

In order to assess whether this heterogeneous subpopulation is conserved in human metastases, we analyzed PDA-derived metastatic TMA cores for Cytokeratin 19, Gstt1 and Ki67. Strikingly, immunofluorescence staining demonstrated intratumoral heterogeneity in metastatic TMA cores, where, similar to the pattern we observed in mouse metastatic tumors, we find that Gstt1-positive cells are mostly (∼80%) non-proliferative (**Figure 3I, 3J**), further validating the presence of a slow-cycling, Gstt1^high^ cell population in human PDA-derived metastases.

### Gstt1^high^ Cells Represent a Slow-Cycling, EMT^high^ Aggressive Metastatic Population

To dissect the function and molecular features of the Gstt1^high^ metastatic subpopulation, we employed CRISPR technology to tag the *Gstt1* endogenous mouse locus to an *mCherry* reporter (**Figure 4A**). Proof of principle *in vitro* analysis and sorting of mCherry populations in PDAC-derived lung metastatic cells generated to express *mCherry* from the *Gstt1* locus demonstrated enriched *mCherry* and *Gstt1* expression in *mCherry^high^* sorted cells (**Figure 4B**). We then performed gene expression profiling of mCherry^low^ and mCherry^high^ populations. To achieve accurate representation of gene expression across Gstt1^high^ and Gstt1^low^ subpopulations as they exist *in vivo*, we first injected GFP-positive PDAC-derived lung metastatic cells (n=3, 1 x 10^5^ cells) expressing endogenous Gstt1-mCherry retro-orbitally into mice to form lung metastases. Once the mice developed macrometastases, entire lungs were dissociated, followed by sorting of mCherry^low^ (bottom 20%) and mCherry^high^ (top 20%) populations gated on GFP to eliminate any contamination from the lung stroma. mCherry^low^ and mCherry^high^ sorted populations were collected for whole genome transcriptome profiling using RNA-Seq (**Figure 4C, 4D**). Unsupervised principal component analysis (PCA) identified distinct clusters of mCherry^high^ and mCherry^low^ populations consistent with enriched Gstt1 expression in mCherry^high^ tumor cells (**Figure S6A, Figure 4B**). Gene expression analysis revealed distinct profiles of differentially expressed genes between mCherry^high^ and mCherry^low^ populations (2FC, FDR 0.05) (**Figure 4D, Table 2**) with biological pathways involved in cell cycle and cell division downregulated in the mCherry^high^ population, further supporting the slow-cycling nature of Gstt1^high^ cells (**Figure 4E**). In addition to cell cycle pathways, gene set enrichment analysis (GSEA) of our mCherry^high^ population independently confirmed an EMT, TGF-β and angiogenesis gene expression signature previously identified as critical features of latent PDA metastases (**Figure S6B**) (*36, 37*). This was confirmed in our mCherry populations via western blot, with an enrichment in vimentin (EMT marker) and a downregulation of E-cadherin in Gstt1^high^/CK19^low^/PCNA^low^ cells (**Figure 4F**). Consistent with functional results and with Gstt1 as a regulator of a slowly proliferating metastatic subpopulation, genome-wide transcriptional profiling on *Gstt1* depleted metastatic cells demonstrated an enrichment in cell cycle, cell division and mitotic microtubule formation pathways (2FC, FDR 0.01) (**Figure S6C**). Complementary to results observed in Gstt1^low^ tumor cells, inhibiting *Gstt1* resulted in an increase in cell cycle genes involved in cell division (*Wee1, Cenpe, Ccnf, Kntc1*) (**Figure S6D**), together pointing to Gstt1 as a novel regulator of metastatic cell proliferation.

**Figure 4.**
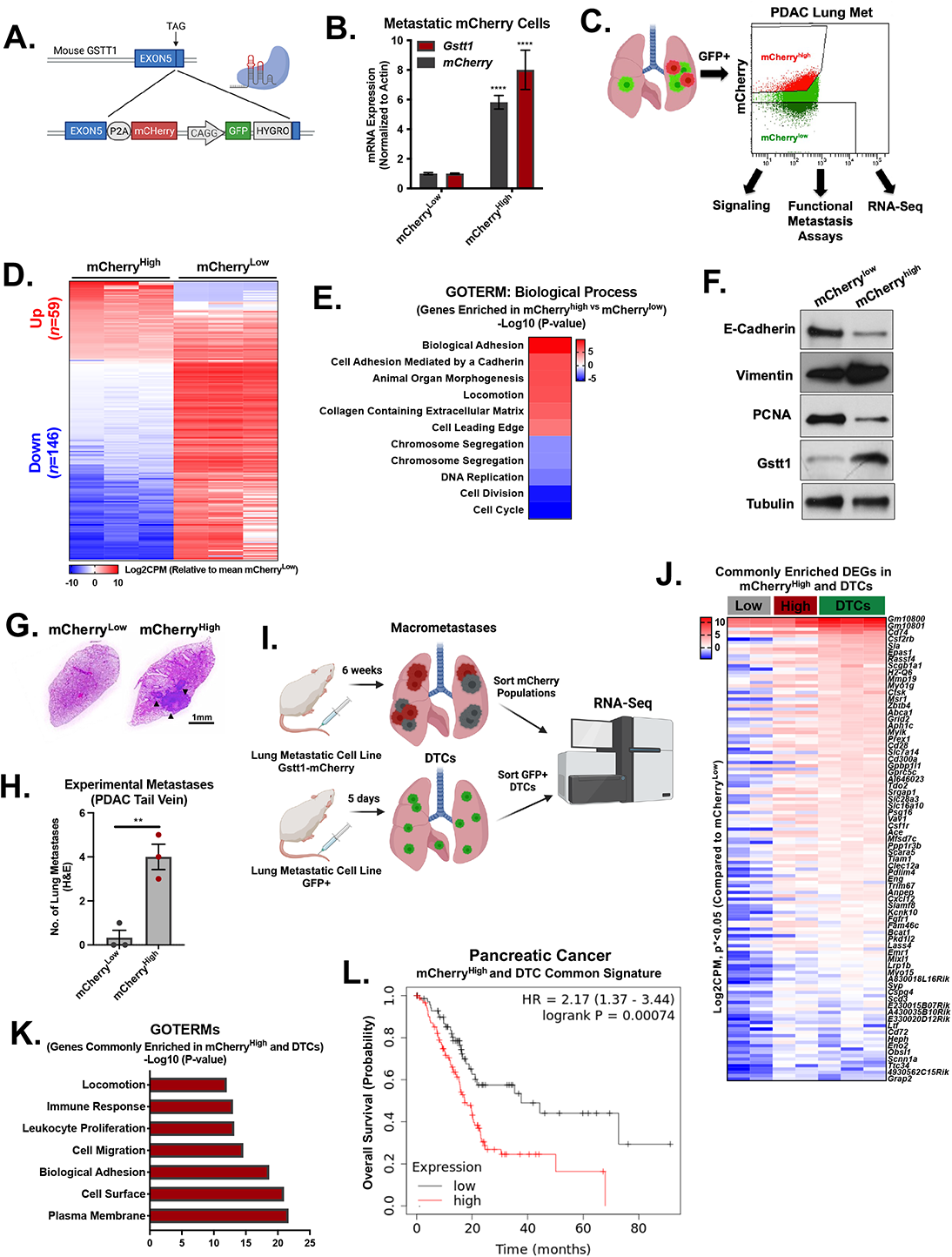
(A) Diagram depicting Cas9 targeting of *mCherry* into the endogenous mouse *Gstt1* locus. (B) qRT-PCR expression levels of *mCherry* and *Gstt1* in lung metastatic cells stably expressing the Gstt1-mCherry construct from three independent FACS experiments. (C) PDAC-derived lung metastatic cells generated to express *GFP+* and endogenously tagged *Gstt1-mCherry* were injected retro-orbitally into SCID mice. Upon evidence clinical signs of metastasis (hunched posture, weight loss), lungs were dissociated, and tumor cells were isolated from the lung stroma using fluorescence activated cell sorting (FACS) (GFP+ gating). Once all tumor cells were gated on GFP+, top (20%) mCherry^high^ and bottom (20%) mCherry^low^ populations were sorted and collected for signaling experiments, functional metastasis assays and RNA-Sequencing. (D) FACS of mCherry populations from *in vivo* tumors were subjected to RNA-Seq. Heatmap depicting differentially expressed genes (DEGs) enriched in the mCherry^high^ population (59 UP, 146 DN) (Log2CPM, Log<2FC, FDR 0.05). (E) DAVID biological pathway analysis demonstrates GOTERM gene signatures enriched and downregulated in mCherry^high^ metastatic cells. (F) Western blot analysis of E-cadherin, Vimentin (EMT), PCNA and Gstt1 in FACS sorted mCherry^high^ and mCherry^low^ populations. (G) mCherry^high^ and mCherry^low^ populations (5 x 10^4^ cells per group) were injected via tail vein into SCID mice to generate experimental metastases to the lung. Mice were sacrificed when demonstrating clinical signs of metastasis (hunched posture, weight loss). Quantification of metastatic burden in H&E stained slides. Data are represented as mean s.d. *t-test* was used to determine statistical significance between groups (p**<0.01). (H) Representative image of H&E stained mCherry lungs from each group. (I) PDAC-derived lung metastatic cells generated to express GFP+ and endogenously tagged Gstt1-mCherry were injected via tail vein into SCID mice (n=3 mice). After 5 days, lungs were dissociated, and tumor cells were isolated from the lung stroma using fluorescence activated cell sorting (FACS) (GFP+ gating). GFP+ tumor cells (DTCs) were subjected to RNA-Seq and analyzed compared to mCherry^low^ and mCherry^high^ macrometastatic populations from (D). (J) Heatmap depicting commonly enriched genes (DEGs) in DTCs and mCherry^high^ population (139 genes) compared to mCherry^low^ populations (Log2CPM, 2FC, p*<0.05). (K) DAVID biological pathway analysis demonstrates GOTERM gene signatures commonly enriched DTCs and in mCherry^high^ macrometastatic cells. (L) Commonly upregulated genes in DTCs and mCherry^high^ populations (<2FC, n=40) were analyzed for impact on overall survival in pancreatic cancer (KMplotter).

**Table 1.** Bulk Tissue RNA Seq DEGs.

**Table 2.** mCherry Hi vs. Low RNA-seq DEGs.

To directly assess the functional significance of the Gstt1^high^ population, we first sorted mCherry populations and performed soft agar assays *in vitro*. mCherry^high^Gstt1^high^ cells were significantly more efficient at forming colonies in 3D (**Figure S7E, S7F**), and importantly, overexpression of *Gstt1* in the Gstt1^low^ cells was sufficient to increase colony numbers to the same level as Gstt1^high^ cells (**Figure S7H, S7I**), further indicating that Gstt1 is required and sufficient for enhancing anchorage independent growth.

Previous reports in head and neck cancer have demonstrated that slow cycling populations function as metastasis-initiating cells (*33*). To functionally test whether the slow growing Gstt1^high^ population is more efficient at promoting metastasis, we injected equal numbers (5 x 10^3^ cells per mouse) of our sorted mCherry^high^ and mCherry^low^ cells via tail vein to generate experimental lung metastases. *mCherry^high^* (*Gstt1^high^*) expressing cells formed significantly more experimental metastases suggesting *Gstt1^high^*cells are endowed with metastasis initiating potential (**Figure 4G, 4H**). Together, these results indicate that Gstt1^high^ cells represent a highly metastatic, slow-cycling population within established metastases with enhanced EMT features.

### Gstt1^high^ Metastases Retain Features of Latent Disseminated Tumor Cells

Since we observe Gstt1^high^ cells in both DTCs and within macrometastases, we asked whether the Gstt1^high^ population in established metastases retains characteristics of latent disseminated tumor cells. To do this, we compared gene expression profiles from our mCherry macrometastatic populations, to DTC RNA profiles. DTCs were generated from experimental metastases, 5 days after tail vein injection and subsequently sorted based on GFP (**Figure 4I**). As we previously demonstrated, we find that almost 100% of DTCs are Gstt1-positive (**Figure 2G**). Expression analysis comparing mCherry^high^ and DTC populations to mCherry^low^ cells demonstrated 139 commonly differentially expressed transcripts (2FC, p*<0.05) with pathways enriched in biological adhesion, cell migration and locomotion (**Figure 4J, 4K, Table 4**). Strikingly, this Gstt1^high^ upregulated transcriptional signature is associated with poor overall survival pancreatic cancer patients (**Figure 4L**), supporting our findings that Gstt1^high^ cells display enhanced metastasis initiating capabilities.

**Table 3.** Control vs. shGstt1 RNA-seq DEGs.

**Table 4.** mCherry Hi vs. DTCs Compared to mCherry Low RNA-seq DEGs.

To determine whether Gstt1^high^ metastatic cells retain dissemination signatures, we applied our transcriptional profiles using an established “early dissemination” gene expression dataset (*20*). We found that mCherry^high^/Gstt1^high^ cells had retained several dissemination markers (*Myo9a*, *Dclre1c*, *Ahnak*, *Ddx46*, *Rpb1*), to similar levels seen in DTCs (**Figure S6E**), which together with cell adhesion and migration gene signatures could explain the enhanced metastasis initiating potential observed in mCherry^high^/Gstt1^high^ cells. This was associated with a decreased proliferation signature in mCherry^high^ cells, with transcript levels of *Ki67*, *Wee1* and *Pcna* comparable to latent DTCs (**Figure S6F**). Altogether, these results demonstrate that the Gstt1^high^ subpopulation within established metastases is a slow-cycling subpopulation that retains latent disseminated cell features and enhanced metastasis initiating potential.

### Gstt1 Glutathionylates Intracellular Fibronectin, a Key Step to Enhance its Secretion and to Promote Gstt1-dependent Metastasis

Although Gstt1 has been found to detoxify xenobiotic compounds (*3,50,52*), its role in metastatic lesions remains unknown. In order to define specific functions for Gstt1 in the context of metastatic tumors, we decided to first investigate interactors and protein substrates of this glutathione transferase. For this purpose, extracts from PDAC-derived liver and lung metastatic cell lines were immunoprecipitated (IP) using both an anti-Gstt1 and a pan-glutathione antibody (anti-GSH) and subjected to mass spectrometry analysis (**Figure S7A, Table 5**). We identified 18 commonly enriched peptides between both Gstt1 and GSH immunoprecipitates (**Figure 5A**). Strikingly, top peptides commonly enriched encode for proteins involved in cytoskeletal, mitotic cell cycle and cell junction pathways (**Figure 5B**). These structural Proteins include Plectin, Fibronectin, Filamin A, Filamin B and Vimentin, however, Fibronectin demonstrated the highest peptide enrichment in *Gstt1* overexpressing cells (**Figure 5C**). Of these, endogenous Fibronectin and Plectin were validated in both Gstt1 and GSH pulldowns from metastatic lysates (**Figure 5D** and **Figure S7B**), suggesting functional significance of this intracellular interaction between Gstt1 and cytoskeletal junction proteins.

**Figure 5.**
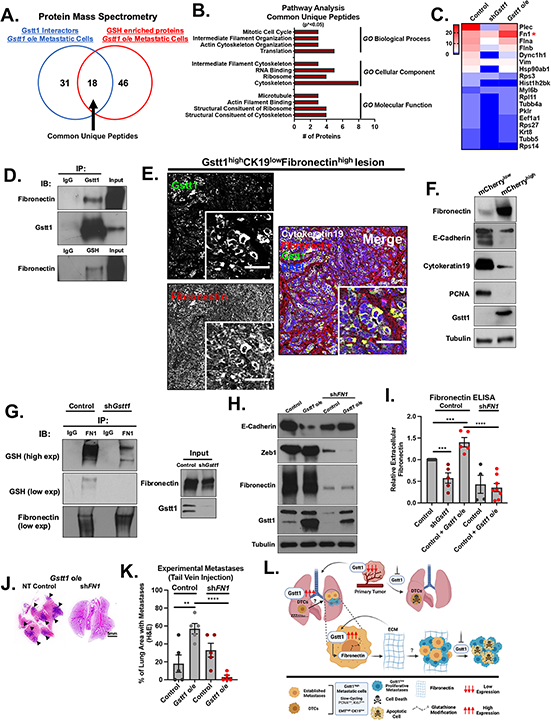
(A) Lung and liver metastatic cell lines (PDAC) were lysed and subjected to immunoprecipitation using a Gstt1 and pan-glutathione (GSH) antibody (whole cell lysate). Pull downs were analyzed for interactors and enriched proteins using unbiased mass spectrometry (M/S). Mass spectrometric analysis identified 18 peptides commonly enriched in both Gstt1 and GSH pulldowns. (B) 18 commonly enriched peptides mainly fell into pathways involved in cytoskeletal, cell junction and structural molecule activity (GO_TERMs). (C) Heatmap depicting number of unique peptides per group as identified by both Gstt1 and GSH pull-down mass spectrometry. Fibronectin (red asterisk), plectin, filamin A, filamin B and vimentin were chosen for further validation. (D) Gstt1 and GSH pull downs in liver metastatic cells were blotted for validation of the interaction with fibronectin. (E) Confocal imaging of immunofluorescence staining of Gstt1, Fibronectin and Cytokeratin 19 (pancreatic cell marker) in CK19^high^Gstt1^low^ liver metastatic lesions. (F) Western blot analysis of Fibronectin, E-cadherin, Cytokeratin 19, PCNA and Gstt1 in FACS sorted mCherry^high^ and mCherry^low^ populations. (G) Liver metastatic cell lines (PDAC) expressing either Control or sh*Gstt1* were lysed and subjected to immunoprecipitation using Fibronectin antibody (whole cell lysate). Fibronectin pull downs in liver metastatic cells were immunoblotted using both fibronectin and pan-GSH antibody. (Right panel) Input controls. (H) Liver metastatic cell lines (PDAC) stably expressing control or i*Gstt1* cDNA were generated to stably express control or sh*FN1.* Cells were subsequently lysed and analyzed via western blot. (I) Media collected from liver metastatic (PDAC) cell lines (2D) stably expressing Control, sh*Gstt1* or i*Gstt1* cDNA were subjected to an ELISA assay for extracellular Fibronectin deposition. Fibronectin levels were normalized to cellular protein content. Data is normalized to Control. *t-test* was used to determine statistical significance between groups (p***<0.005; p****<0.0001). (J) Metastatic cell lines from (H) were injected via tail vein into SCID mice to generate lung metastases. 2 weeks after injection, mice were given doxycycline to induce Gstt1 expression. Mice were sacrificed when demonstrated clinical signs of metastasis (hunched posture, weight loss). (J) Representative H&E images of lung quantification from panel (K). (K) Lung metastatic burden was evaluated using H&E and represented as total lung tissue area versus the metastatic tissue area. Data are represented as mean s.e.m. *t-test* was used to determine statistical significance between groups (p**<0.01, p****<0.0001). (L) Model.

**Table 5.** Mass Spec Gstt1 and GSH pulldowns.

Confocal imaging of liver sections containing multiple metastases identified extracellular Fibronectin fibrils localized in the periphery of metastatic lesions (**Figure 5E** and **Figure S7C**). Surprisingly however, we also detected co-localization of intracellular Fibronectin exclusively within Gstt1^high^ tumor cells (**Figure 5E** and **Figure S7C**). This enrichment of intracellular fibronectin and downregulation of CK19 levels was mirrored in our mCherry^high^/Gstt1^high^ isolated populations (**Figure 5F**). We next assessed whether Gstt1 could be directly glutathione-modifying Fibronectin as previously observed in other ECM proteins (*2*). Indeed, we were able to detect glutathione modification on Fibronectin in metastatic cells, and such modification was almost completely inhibited upon inhibition of *Gstt1* (**Figure 5G**). To assess if Gstt1 enzymatic activity directly modulates glutathionylation of Fibronectin, we performed immunoprecipitation of Fibronectin in the presence of wild-type *Gstt1* and a previously described *Gstt1* catalytically dead mutant, *Gstt1*-R234G (*44, 45*). In the context of wild-type *Gstt1* overexpression we were able to detect a glutathione modification on Fibronectin, however, the modification was absent in the presence of the *Gstt1* catalytically dead mutant (**Figure S7D**). Next, we wanted to determine whether the *Gstt1* catalytically dead mutant could rescue 3D growth defects previously observed in mCherry^low^ cells. As previously mentioned, overexpression of wild-type *Gstt1* was able to rescue soft agar colony growth to levels seen in mCherry^high^ cells, however, the catalytically dead mutant (*Gstt1*-R234G) showed only a partial rescue in 3D growth, suggesting that the glutathione-transferase activity of Gstt1 could be crucial for the acquisition of metastatic traits (**Figure S7E, S7F, S7G**). All together, these results identify Fibronectin as a putative novel substrate of Gstt1 in metastatic cells.

Extracellular matrix (ECM) proteins, such as Fibronectin have been shown to be integral to the maintenance of a supportive metastatic niche (*7,13,34,43*). However, these studies have focused solely on the extracellular deposition of Fibronectin fibrils as critical for maintenance of metastatic phenotypes. The presence of intracellular Fibronectin staining largely restricted to Gstt1-positive metastatic cells (**Figure 5E, 5F** and **Figure S7C**) suggests a biological role for Fibronectin in the aggressive metastatic traits observed in this specific cell population. To begin to identify the functional role of intracellular Fibronectin on Gstt1-induced metastasis, we inhibited Fibronectin in both Gstt1^high^ (*Gstt1* overexpressing and mCherry^high^ sorted cells) and Gstt1^low^ metastatic cell lines. Functionally, *FN1* (Fibronectin gene) depletion demonstrated that mCherry^high^/Gstt1^high^ cells where highly dependent on Fibronectin for soft agar growth (**Figure S7H, S7I**), and depletion of Fibronectin in Gstt1^high^ cells reverted these cells to an epithelial phenotype (**Figure 5H, Figure S7J**), indicating acquisition of less aggressive metastatic characteristics. Of note, inhibition of *FN1* resulted in a reduction in Gstt1 levels (**Figure 5H, S7J**), suggesting a feedback mechanism in which Gstt1 regulates Fibronectin secretion, in turn influencing Gstt1 levels in high expressing metastatic cells. In order to directly assess whether glutathionylation of Fibronectin by Gstt1 regulated its secretion, we measured Fibronectin secretion *in vitro* upon over-expression and knockdown of *Gstt1* using an ELISA assay. Strikingly, knocking down *Gstt1* significantly reduced the secretion of Fibronectin, while overexpressing *Gstt1* increased it, a phenotype rescued upon *FN1* knockdown (**Figure 5I**).

To directly assess whether Gstt1 was acting through Fibronectin to generate metastasis, we injected into mice WT or *Gstt1* overexpressing cells where we inhibited Fibronectin. *FN1* knockdown alone did not reduce the number of macrometastases (**Figure 5J, 5K**) in Gstt1^low^ expressing metastatic cells, suggesting that Fibronectin is not universally required for metastatic cell growth in PDAC. However, *FN1* knockdown completely rescued the increased metastases we observed in *Gstt1* overexpressing cells, highlighting its requirement specifically in the Gstt1^high^ metastatic cell population (**Figure 5J, 5K**) Altogether, our data supports a mechanism whereby Gstt1 directly modifies intracellular Fibronectin to influence EMT and metastatic traits by regulating Fibronectin deposition in the extracellular metastatic tumor microenvironment, in turn identifying Gstt1 as a critical modulator of metastatic disease (**Figure 5L**).

## DISCUSSION

Utilizing mouse models of spontaneous metastasis, we have developed a targeted, functional loss-of-function screen to identify non-genetic factors important for metastatic growth. Although we focused primarily on PDAC, these genes were found to be important for both pancreatic and breast cancer-derived metastases, providing a rationale that these genes may be required across cancer types. Validation in human metastatic PDA studies and their effect on patient survival highlight the specific requirement for metastatic growth. We identify Gstt1 as highly enriched in metastatic lesions and not in primary tumors, acting as a modulator of a subpopulation of slow-cycling metastatic cells. *In vivo* studies demonstrate a specific requirement for *Gstt1* for metastatic growth and dissemination, however dispensable for primary tumor growth. We observed that virtually all DTCs express Gstt1 and that characteristics of slow cycling disseminated cells are retained within the Gstt1^high^ subpopulation of established macrometastatic lesions. This subset of CK19^low^Gstt1^high^ cells appears to be slow cycling, EMT enriched and highly metastatic. Mechanistically, we find that Gstt1 interacts with and glutathione-modifies intracellular Fibronectin, influencing Fibronectin deposition into the metastatic tumor microenvironment critically necessary to drive metastasis. Previous work postulated that the majority of Fibronectin contribution comes from Cancer Associated Fibroblasts (CAFs) (*43*). Our work challenges this notion, indicating that the tumor cells within the metastatic niche appears to play a critical role in the secretion and deposition of Fibronectin into the ECM. Consistent with our results, at least one previous study found that, indeed, Fibronectin contribution from both the tumor cells and the stroma of the primary tumor niche, with implications for enhanced metastasis (*32*). Future studies will define whether Gstt1-dependent glutathionylation of Fibronectin is required for secretion into the metastatic tumor microenvironment.

Glutathione S-transferase theta 1 (Gstt1) is a member of a superfamily of proteins that catalyze the conjugation of glutathione to a variety of hydrophobic and electrophilic molecules. Gstt1 has predominantly been characterized in the liver, where it participates in the detoxification of exogenous xenobiotic compounds (*3,35,44,45*). Interestingly, the *Gstt1* gene is absent in 10-30% of the population, and the null genotype has shown conflicting results in the risk of development of numerous cancers (*3,30*,*35,52*); however, these studies have been descriptive, and do not extend to the context of metastatic disease. Future studies will be required to stratify specifically metastatic patient samples based on the presence of the *Gstt1*-null polymorphism, to determine whether the null allele correlates with survival outcomes.

A recent single-cell lineage study in a mouse model of PDAC revealed an EMT-continuum common to all disseminated metastases, wherein hybrid-EMT states correlate with the most aggressive clones (*47*). Notably, the majority of metastatic clones resided in a single transcriptional cluster. Due to the highly aggressive and rapidly metastasizing nature of their PDAC model system, their data infers that the H1 hybrid-EMT cluster may represent a latent subpopulation of metastatic cells. Interestingly, the majority of our 136 gene ‘metastatic signature’ genes identified within these clusters (n=18) fell into the H1 category (data not shown), with *Gstt1* expression highly enriched in metastatic clones within that cluster. Consistent with our findings in mCherry^high^/Gstt1^high^ populations, the H1 cluster was devoid of proliferative markers (G2M, E2F and mitotic spindle gene sets) and was highly enriched in pathways involved in xenobiotic metabolism, apical surface markers and cell adhesion similar to a previous study which also identified a slow cycling subpopulation of metastasis initiating cells in head and neck cancer (*33*). Additionally, the top differentially expressed genes we found to be commonly enriched in the mCherry^high^/Gstt1^high^ population and downregulated in sh*Gstt1* cells (*Elf3, Spns2*, *C77370*, *1700008I05Rik, Nexmif*), also fall into the H1 EMT cluster, some of which have been shown to be important for metastatic colonization (*54*). Characterization of the H1 cluster, and whether *Gstt1* regulates this cluster in a more slowly progressing model system could provide further insights into mechanisms of dissemination and colonization in pancreatic cancer.

Metastatic colonization, growth and maintenance are strongly influenced by cues from the metastatic tumor microenvironment (*1, 13, 21, 34, 36, 41, 44*) and recent studies have laid the groundwork for targeting the ECM as a mechanism for sensitizing slow growing disseminated tumor cells to chemotherapy (*7,13,16,34,43*). Our work identifies a novel, subpopulation of slow cycling Gstt1^high^/CK19^low^ metastatic cells which express intracellular Fibronectin to promote metastasis. Strikingly, our work points to a feedback loop between Gstt1 modification of intracellular Fibronectin levels and ECM deposition, and in turn Fibronectin signaling regulating Gstt1 expression (**Figure 5H, Figure S7F**). This suggests an important relationship between the tumor microenvironment and cell autonomous mechanisms in governing metastatic heterogeneity. Future studies will be required to determine how *Gstt1* RNA levels are regulated within and across metastatic tumors.

In summary, *Gstt1* has emerged from a loss-of-function shRNA screen as a novel mediator of metastatic disease. We have found that Gstt1 is preferentially upregulated in both mouse and human metastases and functions as a specific driver of metastases, without influencing the growth of primary lesions. Effective treatments for eliminating metastases are limited due to distinct non-genetic heterogeneity from primary tumors and across metastatic tumors as well as the slow cycling nature and lack of detection methods for disseminated cells. The identification of this novel subpopulation required for the maintenance of established metastases provides insight into metastatic heterogeneity and a rationale for the targeting of Gstt1 to treat metastatic disease.

### EXPERIMENTAL PROCEDURES

#### Lead Contact and Materials Availability

Further information and requests for resources and reagents should be directed to and will be fulfilled by the Lead Contact Raul Mostoslavsky.

#### Mouse Models

Mice were housed in pathogen-free animal facilities. All experiments were conducted under protocol 2019N000111 approved by the Subcommittee on Research Animal Care at Massachusetts General Hospital.

The PDAC genetically engineered models (*p48-Cre/p53F/+KrasL/+* (*Sirt6 WT* and *KO*)) have been previously described (*22,41,42*) and were maintained on a mixed 129SV/C57BL/6 background. Mice develop spontaneous pancreatic primary tumors, peritoneal carcinomatosis and distant metastases to the liver and the lung. Mice were sacrificed for all experiments at end point as evidenced by overall poor body condition in accordance with animal welfare guidelines. Poor body condition is characterized as ascites, hunched posture, emaciation and lethargy. Metastatic tissues were considered ‘High Burden’ or ‘Low Burden’ based on the presence of macrometastases or absence of overt metastatic tumors, respectively. All tissues were evaluated for metastases by a pathologist (R.Bronson) at the DF/HCC Research Pathology Core.

The 4T1 breast cancer (BC) metastasis model (*6*) can spontaneously metastasize from the primary tumor in the mammary gland to multiple distant sites preferentially the lung. To generate lung metastases, female BALB/c mice were injected with 1 x10^5^ 4T1 breast cancer cells into the 4^th^ inguinal mammary fat pad. After 4 weeks, mice were sacrificed, and tumors collected for experiments.

For the analysis and prospective isolation of tumor cell populations from both primary tumors and spontaneous metastases, we generated *p48-Cre/p53F/+KrasL/+/ROSA26-LSL-YFP* (*Sirt6* cKO) animals, as well as *Sirt6* WT *p48-Cre/p53F/+KrasL/+/ROSA26-LSL-YFP* animals to specifically isolate YFP^+^ epithelial cells by FACS. 4T1 breast cancer cells were stably transduced with retrovirus containing pMSCV-luc-PGK-Neo-IRES-eGFP to generate GFP expressing cells for sorting experiments.

#### RNA Isolation from Mouse Tumors and RNA Sequencing

Tumor bearing mice were euthanized and both primary tumors and visible metastatic tumors were excised, minced and flash frozen in dry ice. Flash frozen tissues were homogenized using a benchtop homogenizer (Precellys 24, Bertin Instruments) and bulk tissue RNA isolation was conducted using the RNeasy Mini Kit (Qiagen, 74104) according to the manufacturer’s instructions.

cDNA was synthesized and RNA-seq libraries were constructed using the TruSeq RNA Sample Prep kit from Illumina. Libraries were quantified using the Library Quantification kit (Kapa Biosystems, KK4828) and the Bio-Rad CFX96 instrument. Each lane of sequencing was pooled in 6-plex (six samples per lane) with unique barcodes. Pooled libraries were also quantified using the Kapa Biosystems Library Quantification kit (KK4828) and the Bio-Rad CFX96 instrument. These pools were then denatured to 16 pM with 1% PhiX and sequenced on the Illumina HiSeq 2000 instrument, producing approximately 30 million paired-end 50-bp reads per sample.

#### Mouse Tumor Cell Isolation, Fluorescence-Activated Cell Sorting and RNA Sequencing

*p48-Cre/p53F/+KrasL/+/ROSA26-LSL-YFP*(*Sirt6* cKO), as well as *Sirt6* WT *p48-Cre/p53F/+KrasL/+/ROSA26-LSL-YFP* as well as 4T1-eGFP tumor bearing mice were euthanized and entire tissues (primary tumors, livers and lungs) were minced and incubated with 1.3mg/ml collagenase (Sigma-Aldrich, C6079) on a horizontal shaker at 37 °C for 30 min, vortexing every 5 min. Serially filtered and digested tissues using 70-μm and 40-μm strainers (BD Biosciences) were inactivated using 10% FBS in HBSS and centrifuged at 1,200 r.p.m. at 4 °C for 10 min. In order to obtain a single cell suspension, incubate digested tissues with 0.25% Trypsin-EDTA. Cell pellets were resuspended with PBS containing 4% chelated FBS and then transferred into FACS tubes with a 40μm filter. DAPI (4′,6-diamido-2-phenylindole) (3 nM) was added to the cell suspension to negatively select live cells. Proper single-color and negative controls were used in every experiment to optimize gating. Cells were analyzed and sorted using a FACSAria II (BD Biosciences). Obtained data were analyzed by FlowJo.

Sorted cells were collected directly into RNA isolation buffer provided by the following kit. RNA isolation was conducted using the RNAqueous™-Micro Total RNA Isolation Kit (Thermo, AM1931) according to the manufacturer’s instructions. cDNA was synthesized and RNA-seq libraries were constructed using the SMART-Seq v4 Ultra-Low Input RNA kit to produce cDNA (Clontech, 634888). Libraries were quantified using the Library Quantification kit (Kapa Biosystems, KK4828) and the Bio-Rad CFX96 instrument. Each lane of sequencing was pooled in 6-plex (six samples per lane) with unique barcodes. Pooled libraries were also quantified using the Kapa Biosystems Library Quantification kit (KK4828) and the Bio-Rad CFX96 instrument. These pools were then denatured to 16 pM with 1% PhiX and sequenced on the Illumina HiSeq 2000 instrument, producing approximately 30 million paired-end 50-bp reads per sample.

### RNA Sequencing Analysis

STAR aligner was used to map sequencing reads to the mouse reference transcriptome (mm9 assembly) (*4, 12*). Read counts over transcripts were calculated using HTSeq (v.0.6.0) based on a current Ensembl annotation file for NCBI37/mm9 assembly. Differential expression analysis was performed using EdgeR (*40*), and genes were classified as differentially expressed based on cut-offs of at least two folds change and <FDR 0.05 where indicated. Analysis of enriched functional categories among detected genes was performed using DAVID (*49*).

### Cell Lines

To establish mouse pancreatic cancer cell lines from both primary tumors and lung and liver metastases, freshly isolated tissues specimens from *p48-Cre/p53F/+KrasL/+* (*Sirt6 WT* and *KO)* mice were minced with sterile razor blades, digested with trypsin for 30 mins at 37°C, and then resuspended in RPMI 1640 and supplemented with 10% fetal bovine serum and 1% penicillin (100 U/ml)/ streptomycin (100 Ug/ml) (Invitrogen Gibco) and seeded on plates coated with rat tail collagen (BD Biosciences). Cells were passaged by trypsinization. All studies were done on cells cultured for less than ten passages. Cells were cultured in RPMI 1640 supplemented with 10% fetal bovine serum and 1% penicillin (100 U/ml)/ streptomycin (100 Ug/ml) (Invitrogen Gibco). A total of 12 independent cell lines from liver and lung metastases were used for subsequent *in vitro* and *in vivo* experiments. Metastasis-derived human pancreatic cancer cell lines, KP3, KP4 and CFPAC1 were obtained from the Center for Molecular Therapeutics at the Massachusetts General Cancer Center. Cell were cultured in RPMI 1640 supplemented with 10% fetal bovine serum and 1% penicillin (100 U/ml)/streptomycin (100 Ug/ml) (Invitrogen Gibco).

### Targeted shRNA Library Preparation

A library of 470 shRNAs targeting our top 94 mouse identified to be differentially expressed genes between primary tumors and metastases was assembled in the lentiviral vector pLKO.1 and obtained from the Molecular Profiling Laboratory at Massachusetts General Hospital Cancer Center. For each of the 94 gene targets, 5 individual shRNA plasmids were obtained in 96-well format. Transfection-grade DNA was obtained for each individual shRNA plasmid utilizing the Qiagen Plasmid Plus 96 Miniprep Kit (16181, Qiagen).

### shRNA Library Transduction

Individual lentiviruses for the targeted library were prepared as per the 96-well format lentivirus shRNA Broad Institute Genetic Perturbation Platform (https://www.broadinstitute.org/genetic-perturbation-platform).

### Anchorage-Independent shRNA Screen

Cells from liver (PDAC-derived, n=3) and lung (PDAC-derived, n=3; BC-derived, n=3) metastases were plated into 96 well plates (1 x 10^3^ cells per well). Utilizing individual shRNAs for each gene target (94 genes x 5 pooled shRNAs per target), metastases-derived cell lines were infected with 5 pooled shRNA lentiviral particles in a 96-well format to establish stably expressing cells. Stable cell lines were generated via selection using puromycin (5μg ml^−1^) for 48 hours.

After selection, metastatic cells were subjected to a soft agar colony formation assay in a 96-well format (n=9 total cell lines/gene target). Briefly, 96-well plates were coated with 0.6% ‘Bottom Agar’ diluted in media. Stable shRNA expressing metastatic cells were counted, embedded in 0.3% ‘Top Agar’ and plated at a confluency of 1,000 cells per well. After the agarose solidified at room temperature, 100ul of full media was added to the ‘Top Agar’ layer. This feeder layer was replaced every 3 days containing 5μg ml^−1^ puromycin. After two weeks, soft agar colonies were stained using 0.05% p-Iodonitrotetrazolium (INT)-violet overnight (I10406, Sigma), counted and photographed using a Leica DMI 4000b white light microscope. Wells were then ranked by colony number from lowest to highest. Significant differences in soft agar growth were compared to two pLKO.1 scramble shRNA containing wells (p**<0.01, p***<0.005, p****<0.001).

### Human Datasets

To determine differential expression of Top 17 screen hits in human primary vs metastatic PDA, we utilized the Human Metastatic Cancer Database (HMCDB) study derived from McDonald et al, 2017 (*29*). Dataset compares PDA primary tumor derived cells from liver and lung metastases (EXP00257) and is available via GEO Accession (GSE63124). Expression data is represented as “Higher” in metastases vs primary tumors and “Lower” in metastases vs primary tumors or unchanged. *Nr1h3*, *Slc38a4* and *Cd5l* shown to be differentially expressed in human PDA metastases are represented as Log2CPM values and significance was determined using Student’s *t*-test.

To determine whether expression of Top 17 screen hits correlates with Relapse Free Survival in pancreatic cancer we utilized KM plotter (*24*). Dataset includes n=177 patients of all stages. Expression data is represented as “Worse” survival in patients with high overall expression and “Better” survival in patients with higher overall expression.

### Lentivirus Vectors and Stable Cell Lines

Lentivirus were generated by transfecting 293T cell with the indicated vectors (plx304 or pLentiCRISPRv2) and the packaging plasmids deltaVPR and VSVG following the lipofectamine 2000 (Invitrogen) transfection protocol. 293T medium was changed 24 h post-transfection. Virus-containing supernatants were harvested 48 h post-transfection and passed throught a 0.45 um filter. Cells were infected with equal ratio of supernatant containing viral particles to cell culture medium and 8 ug/ml polybrene. Stable cell lines for shRNA knockdowns were generated by infection with the lentiviral vector pLKO.1-puro carrying shRNA sequence for scrambled (Addgene, Cambridge, MA, USA), mouse *Gstt1* (sh#1CACAACTCACAGTTCACAATT;sh#2 CGCCATTTATATCTTCGCCAA;sh#3CCTGTGGCATAAGGTGATGTT;sh#4CCATTTATATCTT CGCCAAGA;sh#5CCTCATCATAAAGCAGAAGCT),human *Gstt1* (sh#1 GCTTGCTTAAGACTTGCCCAA; sh#2CTTTGCCAAGAAGAACGACAT) or mouse *FN1* (sh#5GCCAGTTTCCATCAATTATAA).

To generate lentiviral vectors for doxycycline-inducible overexpression of wild-type mouse *Gstt1*, we replaced the CAS9 sequence in the PCW-CAS9 vector (#50661, Addgene) by the 3X FLAG tagged CDS of TV1 of murine *Gstt1* through Gibson cloning. The *Gstt1* catalytically dead mutant was generated by mutating the 234 amino acid (R-Arginine) to a Glycine (G) using Gibson cloning and the wild-type PCW-FLAG-*Gstt1* as a template (*44, 45*). Stable cell lines were generated by puromycin selection. Cell lines were treated with 1 μg ml^−1^ doxycycline for a minimum of three days to achieve overexpression.

Stable doxycycline-inducible Cas9 cell lines were generated by infection with the PCW-Cas9 vector, followed by puromycin selection and infection with Lenti-sgRNA (104993, Addgene) containing a guide targeting exon 3 of murine Gstt1 (GAACCTTATACTTGTGTGCC). Cells were selected with blasticidin containing media 48 hours after infection. Cell lines were treated with 1 μg ml^−1^ doxycycline for a minimum of three days to achieve KO.

### Tumor Sphere Assay

Metastatic-derived cell lines were plated as single-cell suspension (1x 10^3^ and 1 x 10^4^ cells per well) in ultralow attachment 6-well plates (Corning) and grown in DMEM/F12 medium (serum free) supplemented with 20 μl ml^−1^ B27 (Invitrogen), 20 ng ml^−1^ EGF (Peprotech) and 20 ng ml^−1^ bFGF (Peprotech). Spheres were spun down, and fresh media was added every 3 days. Tumor spheres were counted and photographed at day 10 using using a Leica DMI4000b white light microscope. Tumor sphere assays were performed in triplicate and are represented as mean ± s.d. between three independent experiments.

### Real-Time RTPCR Analysis

Total RNA was extracted with the RNeasy Mini Kit (Qiagen, 74104) as described by the manufacturer. For cDNA synthesis, 1μg of total RNA was reverse-transcribed using the QuantiTect Reverse Transcription Kit (Qiagen). Real-time PCR was run in duplicate using SYBR green master mix (Roche), following the manufacturer’s instructions, with the exception that the final volume was 12.5 μl of SYBR green reaction mix. Real-time monitoring of PCR amplification was performed using the LightCycler 480 detection system (Roche). Data were expressed as relative mRNA levels normalized to the β-actin expression level in each sample and are represented as mean ± s.d. between at least three independent experiments unless otherwise indicated in the figure legend. The primer sequence for mouse *Gstt1* (For-GTTCTGGAGCTGTACCTGGATC; Rev-AGGAACCTTATACTTGTGTGCC), *mCherry* (For-CATCCCCGACTACTTGAAGC; Rev-CCCATGGTCTTCTTCTGCAT) and mouse *β-actin* (For-ACTATTGGCAACGAGCGGTTC; Rev-AAGGAAGGCTGGAAAAGAGCC), primer sequences for mouse *Fgfr3, Nr1h3, Slc38a4, Itga8, Cd5l* and *Ki67* are available upon request.

### Anchorage-Independent Growth Assays

#### Soft Agar Colony Formation

A mixture of 5,000 cells in assay medium and 0.3% agarose was seeded on to a solidified bed of 0.6% agarose on six-well plates. The plates were allowed to solidify at 4°C and incubated at 37°C. The cultures were fed once a week with standard growth medium. Cells were incubated for 21 days and then stained with 500 μl of 0.05% p-Iodonitrotetrazolium (INT)-violet overnight. Colonies (>50 μm in diameter) were counted using a Leica DMI 4000b white light microscope. Tumor sphere assays were performed in triplicate and are represented as mean ± s.d. between three independent experiments.

### Metastasis Studies

#### Breast Cancer Model

##### Validation of shRNA Target Genes

Cell lines derived from 4T1 metastatic lung tumors (see above) were stably transduced with pLKO.1 vectors (Control and pooled shRNAs targeting *Gstt1*), harvested by trypsinization, washed twice in PBS, resuspended at 1 x 10^4^ in 100ul PBS and injected into the venus sinus of BALB/c mice (Jackson Laboratory) to establish lung metastases. Mice were sacrificed for all experiments at end point as evidenced by overall poor body condition in accordance with animal welfare guidelines. Metastatic lesions were confirmed by histological analysis by a pathologist (R.Bronson) at the DF/HCC Research Pathology Core.

##### Primary Tumor Growth Spontaneous Metastases

Cell lines derived from 4T1 primary tumors (see above) were stably transduced with pLKO.1 vectors (Control and pooled shRNAs targeting *Gstt1*), harvested by trypsinization, washed twice in PBS, resuspended at 1 x 10^4^ in 90ul PBS and 10% Matrigel (Corning) and injected into the 4^th^ inguinal mammary fat pad of BALB/c mice (Jackson Laboratory) to establish mammary tumors. Primary tumor growth was monitored weekly by taking measurements of tumor length (L) and width (W). Tumor volume was calculated using the formula pLW^2^/2. Mice were sacrificed when tumors reached 600 mm^3^ as per institutional animal use guidelines. Metastatic lesions in the lungs were confirmed by histological analysis by a pathologist (R.Bronson) at the DF/HCC Research Pathology Core.

For inducible Gstt1 KO experiments, 4T1 cells were first stably transduced with pMSCV-luc-PGK-Neo-IRES-eGFP. This was followed by transduction with iCas9 and guide expressing vectors (Control and sg*Gstt1*), harvesting by trypsinization, washing twice in PBS, resuspension at 1 x 10^4^ in 90ul PBS and 10% Matrigel (Corning), and injected into the 4^th^ inguinal mammary fat pad of BALB/c mice (Jackson Laboratory) to establish mammary tumors. Upon injection, doxycycline (200 μg ml^−1^) was administered in the drinking water of all experimental mice and was replaced every week. Beginning day 7 post-injection, mice were subjected to bioluminescence imaging once a week. Under isoflurane anesthesia, 300 μl of d-luciferin (15 mg ml^−1^) was injected intraperitoneally, and the mice were imaged every 5 min after injection with a 0.5 s to 60 s exposure time on an Ami X imaging system (Spectral Instruments Imaging) until the total flux and maximal radiance peaked. Total flux (photons/s) and maximal radiance (photons/s/cm^2^/sr) were measured by Spectral Ami X (Spectral Instruments Imaging). A region of interest (ROI) was drawn around the mammary fat pad and lung region for each mouse as well as a background ROI outside of the mice, which was subtracted from the total flux and maximum radiance for each mouse. Once the tumors reached 350 mm^3^, the primary tumor was surgically removed under isoflurane anesthesia (*11*). Surgical procedures were performed in the aseptic hood of the pathogen-free facility and mice were singly housed to aid in wound healing. Following primary tumor removal, mice were continuously measured for lung metastases using bioluminescence imaging once a week until reaching end point, or poor body condition in accordance with animal welfare guidelines. Metastatic lesions were confirmed by histological analysis by a pathologist (R.Bronson) at the DF/HCC Research Pathology Core.

##### Metastatic Rescue Experiments

Cell lines derived from 4T1 primary tumors (see above) were stably transduced with pMSCV-luc-PGK-Neo-IRES-eGFP. This was followed by transduction with inducible Gstt1 overexpression and shControl or sh*Gstt1* expressing vectors, harvesting by trypsinization, washing twice in PBS, resuspension at 1 x 10^4^ in 90ul PBS and 10% Matrigel (Corning), and injected into the 4^th^ inguinal mammary fat pad of BALB/c mice (Jackson Laboratory) to establish mammary tumors. Upon injection, doxycycline (200 μg ml^−1^) was administered in the drinking water of all experimental mice and was replaced every week. Beginning day 7 post-injection, mice were subjected to bioluminescence imaging once a week, as described above. Primary tumor growth was monitored weekly by taking measurements of tumor length (L) and width (W). Tumor volume was calculated using the formula pLW^2^/2. Once the tumors reached 350 mm^3^, the primary tumor was surgically removed under isoflurane anesthesia. Surgical procedures were performed in the aseptic hood of the pathogen-free facility and mice were singly housed to aid in wound healing. Following primary tumor removal, mice were continuously measured for lung and disseminated metastases using bioluminescence imaging once a week until reaching end point, or poor body condition in accordance with animal welfare guidelines. Metastatic lesions were confirmed by histological analysis by a pathologist (R.Bronson) at the DF/HCC Research Pathology Core.

#### Pancreatic Cancer Model

##### Validation of shRNA Target Genes

Cell lines derived from *p48-Cre/p53F/+KrasL/+ (Sirt6 WT and KO)* metastatic lung tumors (see above) were stably transduced with pLKO.1 vectors (Control and pooled shRNAs targeting *Gstt1*), harvested by trypsinization, washed twice in PBS, resuspended at 1 x 10^4^ in 100ul PBS and injected into the venus sinus or tail vein of NOD.CB17-Prkdcscid/J mice (Jackson Laboratory) to establish lung metastases. Mice were sacrificed for all experiments at end point as evidenced by overall poor body condition in accordance with animal welfare guidelines. Metastatic lesions were confirmed by histological analysis by a pathologist (R.Bronson) at the DF/HCC Research Pathology Core.

##### Primary Tumor Growth

For PDAC orthotopic injections, cell lines derived from primary tumors from *p48-Cre/p53F/+KrasL/+ (Sirt6 WT and KO)* mice were stably transduced with pLKO.1 vectors (Control and individual shRNA targeting *Gstt1*), harvested by trypsinization, washed twice in PBS, and resuspended at 1 x 10^4^ in 10ul PBS. Mice were anesthetized using isoflurane and cells were surgically injected into the head of the pancreas of NOD.CB17-Prkdcscid/J mice (Jackson Laboratory). Surgical procedures were performed in the aseptic hood of the pathogen-free facility and mice were singly housed to aid in wound healing. Mice were monitored weekly for evidence of poor body condition (ascites, weight loss, hunched posture) and then euthanized as per animal welfare guidelines. Primary tumor volume was calculated using the formula pLW^2^/2. Metastatic lesions in the lungs were confirmed by histological analysis by a pathologist (R.Bronson) at the DF/HCC Research Pathology Core.

##### Metastatic Regression Experiments

Cell lines derived from *p48-Cre/p53F/+KrasL/+ (Sirt6 KO)* metastatic lung tumors (see above) were stably transduced with pMSCV-luc-PGK-Neo-IRES-eGFP. This was followed by transduction with iCas9 and guide expressing vectors (Control and sg*Gstt1*), harvesting by trypsinization, washing twice in PBS, resuspension at 1 x 10^4^ in 100ul PBS and injection into the venus sinus of NOD.CB17-Prkdcscid/J mice (Jackson Laboratory) to establish lung metastases. Beginning day 14 post-injection, all mice were subject to bioluminescence imaging once a week, as described above. Once lung metastatic tumors were detected, doxycycline (200 μg ml^−1^) was administered in the drinking water of all experimental mice and replaced every week. Mice were continuously measured for lung metastases using bioluminescence imaging once a week until reaching end point, or poor body condition in accordance with animal welfare guidelines. At end point, mice were imaged, euthanized and lungs were subjected to *ex vivo* bioluminescence imaging. Metastatic lesions were confirmed by histological analysis by a pathologist (R.Bronson) at the DF/HCC Research Pathology Core.

##### DTC Experiments

Cell lines derived from *p48-Cre/p53F/+KrasL/+ (Sirt6 WT and KO)* metastatic lung tumors (see above) were stably transduced with pLKO.1 vectors (Control and sh*Gstt1*) or iPCW-FLAG-*Gstt1* (treated with doxycycline 5 days prior to injection) were harvested by trypsinization, washed twice in PBS, resuspended at 1 x 10^4^ in 100ul PBS and injected into the tail vein of NOD.CB17-Prkdcscid/J mice (Jackson Laboratory) to establish lung metastases. Beginning day 3 post-injection, all mice were subject to bioluminescence imaging once a week for two weeks. Mice were sacrificed after two weeks, before bioluminescent signal was detected for DTC quantification.

##### Immunoblotting

Cell and tissue lysates were prepared in RIPA lysis buffer (150 mM NaCl, 1% NP40, 0.5% DOC, 50 mM Tris-HCl at pH 8, 0.1% SDS, 10% glycerol, 5 mM EDTA, 20 mM NaF and 1 mM Na3VO4) supplemented with 1 mg/ml each of pepstatin, leupeptin, aprotinin and 200mg/ml phenyl-methyl-sulfonyl-fluoride (PMSF). Lysates were sonicated and cleared by centrifugation at 16000g for 20min at 4°C and analyzed by SDS-polyacrylamide gel electrophoresis (PAGE) and autoradiography. Proteins were analyzed by immunoblotting using the following primary antibodies: anti-Gstt1 (Abcam, ab175418), β-actin (Sigma-Aldrich, A5316), Tubulin (Sigma-Aldrich, T6199), E-Cadherin (BD, 610181), Vimentin (BD, 550513), PCNA (PC10) (Santa Cruz, sc-56), Fibronectin (Abcam, ab199056), GSH (D8) (Santa Cruz, 52399).

##### Histology and Immunostaining

Mouse primary tumors, liver and lungs were harvested, submitted for histological examination and analyzed in a blinded fashion by a pathologist (R. Bronson) at the DF/HCC Research Pathology Core. Tissue samples were fixed overnight in 10% buffered formalin, and then embedded in paraffin and sectioned at a thickness of 5 μm by the DF/ HCC Research Pathology Core. H&E staining was performed using standard methods.

For tissue immunofluorescence, deparaffinization, rehydration and antigen retrieval were performed in unstained slides with Trilogy solution (Cell Marque). Sections were blocked with 5% goat serum (Cell Signaling), 1% BSA (Sigma) and 0.2% gelatin (Sigma) in PBS with 0.1% Triton-X for 1h at RT, and then incubated overnight with primary antibodies at 4 °C. The following primary antibodies and their dilutions were used: anti-Gstt1 (Abcam, ab175418, 1:200 for mouse and human tissues), anti-TromaIII (DSHB, 1:500, for mouse tissues), anti-Cytokeratin (Dako, M351529-2, 1:500, for human tissues), Cytokeratin19-555 (Abcam, 203444, 1:500, for human tissues), anti-PCNA (PC10) (Santa Cruz, sc-56, 1:200, for mouse tissues), anti-Ki67 (Abcam, 1:200, for human tissues), and anti-Fibronectin (Santa Cruz, 271098, 1:200, for mouse and human tissues). Slides were kept in dark containers. Samples were washed three times for 10 min each in PBST and incubated with secondary antibodies for 2h at RT. The following secondary antibodies and their dilutions were used: anti-rabbit, anti-mouse and anti-rat conjugated to Alexa Fluor 488 (Molecular Probe; 1:500), Alexa Fluor 555 (Molecular Probe; 1:500), Alexa Fluor 647 (Molecular Probe; 1:500). Stained slides were mounted using Prolong Gold Antifade mounting reagent (Vector lab, H-1200) containing DAPI for nuclei staining. Pictures were obtained using a Nikon Eclipse Ni-U fluorescence microscope or Leica SP8 Confocal Microscope. Quantification of positive and negative cells was performed manually using ImageJ. Human PDA TMA slide (USA Biomax, PA-2081c) was scanned on the Vectra Polaris scanning microscope and analyzed by the MGH Cancer Center Translational Imaging Core. Average intensity was measured per channel and data is represented as ratio of double positive cells per core (CK19+ and Gstt1+).

##### Label-Retaining Membrane Dye Experiments

PDAC-derived lung metastatic cell lines were labeled with CellTracker Membrane Dye CM-DiI (Invitrogen) as per manufacturer’s instructions and as previously described (*25, 33*). CM-Dil+ and CM-DiI-negative populations were isolated using fluorescence-activated cell sorting (FACS) at indicated timepoints. Representative flow sorting scheme in Fig 4G. Cell populations were collected and analyzed for *Gstt1* and *Ki67* mRNA expression using qRT-PCR and represented as expression relative to Day 0.

### Generation of Gstt1-mCherry Cell Line, FACS and RNA Sequencing

#### Endogenous tagging of murine Gstt1

To create Gstt1-mCherry tagged cell lines, we generated an integration donor made of two 750 bp homology arms flanking the stop codon of murine GSTT1 followed by a selection cassette containing GFP and hygromycin. A CRISPR guide targeting the 150 bp region around the stop codon (G TGCATGGTACCAGCGAGTGG) was cloned into the PX330 vector (#110403, Addgene). 1 x 10^6^ lung metastatic derived PDAC cells were then nucleoporated using a Nucleofector 2b Device (AAB-1001, Addgene) using the Amaxa Cell Line Nucleofector Kit R (VCA-1001, Lonza) and plated into a 10 cm tissue culture dish. After two days in culture, cells were treated 500 ug of hygromycin to select for positive clones. After selection, remaining resistant colonies were picked, expanded, and screened for correct integration by PCR by using the primer pairs #1: For-CAAACCAGGTCTCCATCTGCGTGTCTTG;RevGAACTCCTTGATGATGGCGAATCCTGGTCCG) and Primer pair #2: For-TTCAAGAATGCATGCGTCAATTTTACGCAGACTATCTTTCTAGGG; Rev-TCTCCCCGTCCCCACTCCCAACTACAG.

#### Flow Sorting of mCherry Populations

Lung metastatic derived cell line generated to express mCherry from the endogenous Gstt1 locus (above) was cultured under 2D conditions, trypsinized and filtered through 40µm mesh filters. DAPI (4′,6-diamido-2-phenylindole) (3 nm) was added to the cell suspension to negatively select live cells. mCherry^high^ and mCherry^low^ populations were gated on live populations as well as GFP+. For *in vitro* and *in vivo* assays, top 15-20% and bottom 15-20% were chosen as mCherry^high^ and mCherry^low^ populations, respectively. Proper single-color and negative controls were used in every experiment to optimize gating. Cells were analyzed and sorted using a FACSAria II (BD Biosciences). Obtained data were analyzed by FlowJo. An exemplary gating strategy is described in Fig. 4B. Cells were then plated in soft agar conditions, for tail vein injections (5 x 10^4^ cells) per population per mouse, for western blot analysis and for RNA-Sequencing described below. Data represents a minimum of three technical replicates from three independent sorting experiments.

#### RNA Sequencing on mCherry Populations

Lung metastatic derived cell line generated to express mCherry from the endogenous Gstt1 locus (above) was cultured under 2D conditions, followed by harvesting by trypsinization, washing twice in PBS, resuspension at 1 x 10^4^ in 100ul PBS and injection into the tail vein of NOD.CB17-Prkdcscid/J mice (Jackson Laboratory) to establish lung metastases. Upon evidence of clinical metastasis, mice were euthanized, lung tissue was collected, digested and sorted as described above. mCherry^high^ and mCherry^low^ populations were gated on live populations as well as GFP+. For *in vivo* assays, top 20% and bottom 20% were chosen as mCherry^high^ and mCherry^low^ populations, respectively. Sorted cells were collected directly into RNA isolation buffer provided by the following kit. RNA isolation was conducted using the RNAqueous-Micro Total RNA Isolation Kit (Thermo, AM1931) according to the manufacturer’s instructions. Library construction, RNA Sequencing and analysis were done as described above. In order to obtain a common gene signature, differentially expressed gene cutoff from RNA Sequencing was two folds change, <FDR 0.05. Data represents a minimum of three technical replicates from three independent mice and sorting experiments.

For DTC populations, lung metastatic derived cell line generated to express GFP+ and mCherry from the endogenous Gstt1 locus (above) was cultured under 2D conditions, followed by harvesting by trypsinization, washing twice in PBS, resuspension at 1 x 10^4^ in 100ul PBS and injection into the tail vein of NOD.CB17-Prkdcscid/J mice (Jackson Laboratory) to establish lung metastases. After 5 days, mice were euthanized, lung tissue was collected, digested and sorted as described above. Tumor cells were gated on live populations as well as GFP. GFP+ cells were sorted from 3 independent mice and collected as described above for RNA-Seq analysis.

#### Human Samples

Tumor specimens derived from primary tumors and metastatic lesions from cadavers of rapid autopsy patients analyzed in this study were collected following institutional review board (IRB) approval. The collection and use of de-identified cadaver tissues was reviewed and approved by the Dana-Farber/Massachusetts General Hospital (DF/MGH) IRB (protocol no. 13-416). Freshly isolated specimens from pancreatic cancer primary and metastatic sites were collected and divided into two parts: a) stored in formalin for making FFPE tissue blocks (for H&E and Immunofluorescence); b) bulk tissue homogenization for protein lysates. For this study, numerous primary tumor (if available) and metastatic samples were collected from four individual patients. PDA Tissue Microarray was obtained from USA Biomax (PA-2081c) as an FFPE slide with a total of 192 cores including n=68 primary PDA tumor and n=14 metastatic cores.

#### RNA Isolation from Mouse Metastatic Cell Lines and RNA Sequencing

Cell lines derived from *p48-Cre/p53F/+KrasL/+ (Sirt6 WT and KO)* liver and lung metastases (see above) and grown in 2D culture were stably transduced with pLKO.1 vectors (Control and sh*Gstt1* #1, sh*Gstt1* #2). Cells were collected and RNA isolation was conducted using the RNeasy Mini Kit (Qiagen, 74104) according to the manufacturer’s instructions. Library construction, RNA Sequencing and analysis were done as described above. In order to obtain a common gene signature, differentially expressed gene cutoff from RNA Sequencing was two folds change, <FDR 0.05.

#### Immunoprecipitation (IP) and Mass Spectrometry

PDAC liver metastatic derived cell lines were cultured under 2D conditions as described above. To achieve overexpression of Gstt1, cells were treated with 1 μg ml^−1^ doxycycline for 7 days. Cell lysates were prepared in RIPA lysis buffer, supplemented with protease inhibitors as described above. Lysates were sonicated and cleared by centrifugation at 16000g for 20min at 4°C and subsequently subjected to immunoprecipitation using magnetic Dynabeads Protein-G (Invitrogen, 10007D) immunoprecipitation kits. Prior to immunoprecipitation, magnetic beads were conjugated with specific primary antibodies (Gstt1, Abcam, ab199337; Fibronectin Abcam, ab199056; GSH (D8) Santa Cruz, 52399; Normal Rabbit IgG, Cell Signaling, 2729S). Samples were incubated at 4°C with constant rotation overnight in IP buffer (RIPA, 0.25 mM PMSF plus protease inhibitors cocktail tablets (Sigma-Aldrich, 4693116001)). Magnetic Dynabeads containing immunoprecipitated samples were then washed twice with IP buffer and 3 times with washing buffer supplied in the Dynabead kit. IPs were run on a 4-20% SDS-PAGE gel for 5 minutes before fixing the gel with Coomassie (ProtoBlue Safe, Scientific Labs). Coomassie stained Gstt1 and GSH IPs were submitted to the Taplin Biological Mass Spectrometry Facility at Harvard Medical School for peptide identification. Top enriched peptide interactors commonly identified in Gstt1 and GSH pulldowns were validated using individual IPs.

#### Fibronectin Deposition ELISA

PDAC liver metastatic derived cell lines stably expressing an inducible Gstt1 overexpression construct and sh*Gstt1* were cultured under 2D conditions as described above. To achieve overexpression of Gstt1, cells were treated with 1 μg ml^−1^ doxycycline for 7 days. After 7 days, 1ml of media was collected from all cell conditions. ELISA on collected media was performed as per manufacturer’s instructions (ab108849, Abcam) and normalized to protein concentration for each sample. Experiments were performed in triplicate.

#### Statistical Analysis

All values are expressed as mean ± SD or S.E.M.. Statistical significance was determined using unpaired t-test using Prism version 8 and shown in the Figures whenever applicable. Differences were considered statistically significant at a value of p < 0.05 unless otherwise noted. (* indicates *P* < 0.05; **, *P* < 0.01; ***, *P* < 0.001; ****, *P* < 0.0001 unless otherwise indicated; ns, not significant).

### Data and Software availability

#### GEO Accession numbers

All the raw datasets for the different sequencing experiments will be deposited in NCBI.

## ACKNOWLEDGEMENTS

We would like to thank all the members of the Mostoslavsky lab for critical discussions throughout the years and editing of the manuscript especially Thomas L. Clarke. We would also like to thank Dr. Nabeel Bardeesy for experimental discussions and critical evaluation of the manuscript. R.M. is the Laurel Schwartz Endowed Chair in Oncology. This work is supported by NIH grants (R01CA235412 and R01GM128448) to R.M, and an ACS Postdoctoral Fellowship and NIH grant (K99CA252600-01) to C.M.F. We would also like to thank the members of the HSCI-CRM Flow Cytometry Facility at Massachusetts General Hospital, Dr. Roderick Bronson at the Rodent Histopathology Core at Harvard Medical School, the NextGen Sequencing Core at Massachusetts General Hospital and the Massachusetts General Hospital Translational Imaging Core for their technical expertise.

## Authors Contributions

C.M.F. conducted experiments, wrote the manuscript, designed and interpreted most experiments. R.B., H.C., T.B. and E.R.H. conducted and assisted with experiments. L-P.W., M.C. and R.S. provided all the computational analysis for transcriptomic experiments. G.W. performed mouse bioluminescence imaging. S.K. generated the pancreatic cancer mouse model for both bulk tissue RNA-Seq and cell line generation. D.E.M. and D. J. provided access to rapid autopsy samples and clinical expertise. E.R. and I.M. evaluated pancreatic cancer datasets for bioinformatic analysis of gene expression and survival. R.M. designed experiments, interpreted the data, and wrote and edited the manuscript.

