## Supplemental Figures for "The glutathione S-transferase Gstt1 is a robust driver of survival and dissemination in metastases"

#### SUPPLEMENTAL FIGURE LEGENDS

**Supplemental Figure 1.** (A) Schematic depicting PDAC mouse models used for lineage-traced YFP/GFP+ metastatic cells. Lineage-marked YFP/GFP+ expressing primary, liver and lung metastatic tumor cells were isolated via fluorescence activated cell sorting (FACS) and subjected to RNA-Seq analysis. (B) Expression of Top 15 hits in primary matched GFP+ sorted cells (Log2CPM, no FDR) (significant differential expression between metastases and primary tumors displayed with red asterisk, p*<0.05). (C) qRT-PCR for top 6 hits in primary and matched metastatic PDAC cell lines used for shRNA screen. Data represented as metastatic cell line mRNA expression relative to primary tumor derived cells. Data are represented as mean s.e.m. *t-test* was used to determine statistical significance between groups (p*<0.05; p**<0.01). (D) qRT-PCR for individual shRNA knockdown of top 6 hits in metastatic PDAC cell lines used for validation experiments of screen hits. (E) Brightfield images of INT-violet stained soft agar colonies from liver metastatic cell lines using individual shRNAs validating top 6 screen hits. (F) Number of soft agar colonies per well for each gene target (n=3 cell lines used per shRNA). Data are represented as mean s.e.m. *t-test* was used to determine statistical significance between groups (p*<0.05; p**<0.01, p***<0.005). (G) Differential expression of *Nr1h3*, *Slc38a4* and *Cd5l* in human PDA metastases vs primary tumors in the McDonald et al, 2017 dataset (n=16 primary, liver and lung metastatic samples) EXP00257 on the Human Metastatic Cancer Database. *t-test* was used to determine statistical significance between groups. (H) Tumor sphere growth of *Gstt1* depleted metastatic cell lines. Representative bright-field image (2.5X). (I) Represented as number of tumor spheres per well. The experiment was performed in triplicate with at least 4 independent metastatic-derived cell lines (biological replicates) each. Data are represented as mean s.e.m. *t-test* was used to determine statistical significance between groups (p*<0.05; p**<0.01, p***<0.005). (J) 2D growth curve in metastatic cell lines in response to control or two independent shRNAs targeting *Gstt1*. The experiment was performed in triplicate with three replicates each. Data are represented as mean s.e.m. *t-test* was used to determine statistical significance between groups. (K) Lung metastatic (PDAC) cell lines stably expressing Control, sh*Gstt1* or doxycycline inducible *Gstt1* cDNA were analyzed by western blot for Gstt1 expression. (L) Western blot depicting BC-derived primary and lung metastatic cell lines stably expressing pooled *Gstt1* shRNAs. (M) *In vivo* validation of Gstt1 knockdown demonstrates reduced metastatic burden experimental 4T1 cells (n=5 mice per group). (N) Primary tumor growth curves of orthotopically injected 4T1 cells into the mammary fat pad (n=6 mice per group). (O) Spontaneous metastases derived from 4T1 orthotopic injections in the mammary fat pad (n=6 mice per group) (P) Representative H&E images from spontaneous lung metastases in (N). *In vivo* experiments are represented as mean s.e.m. *t-test* was used to determine statistical significance between groups (p*<0.05; p***<0.005).

**Supplemental Figure 2.** (A) Immunofluorescence staining of Gstt1 and Cytokeratin 19 (pancreatic cell marker) in PDAC-derived primary tumor, liver and lung metastases. (B) Quantification of immunofluorescence staining of Gstt1 populations in all CK19+ primary and metastatic lesions. Data represented as percent of CK19+ cells. Quantification represents the average of 10 fields from a minimum of 5 individual metastatic lesions from n=3 mice. Data are represented as mean s.d. *t-test* was used to determine statistical significance between groups (p****<0.0001). (C) Representative immunofluorescence images of Gstt1 and pan-Cytokeratin (tumor cell marker) in PDA-derived primary matched liver metastases from 3 independent rapid autopsy patients. (D) Western blot analysis of Gstt1 levels in multiple primary (Primary PDA) and matched metastatic tissues (Omental, Lymph Node, Liver and Lung Metastases) derived from two rapid autopsy patients with pancreatic cancer (PDA#1 and PDA #2). (E) Western blot analysis of Gstt1 levels in two metastatic derived human cell lines (KP3 and CFPAC1) stably expressing control or two individual short hairpins targeting *Gstt1*. (F) Quantification of soft agar colony growth in liver metastasis-derived pancreatic cancer cell lines (KP3 and CFPAC1 expressing either control or short hairpins targeting *Gstt1*. The experiment was performed in triplicate with three replicates each. Data are represented as mean s.e.m. *t-test* was used to determine statistical significance between groups (p***<0.005, p****<0.0001).

**Supplemental Figure 3.** (A) Western blot depicting Gstt1 levels in iCas9 + Control or iCas9 + sg*Gstt1* expressing 4T1 cells with and without doxycycline. Balb/c mice were treated with doxycycline upon orthotopic injection of iCas9 + Control or iCas9 + sg*Gstt1* expressing 4T1 cells into the mammary fat pad (n=5 mice per group). Mice were monitored for primary tumor growth via bioluminescence imaging and primary tumors were surgically removed when the tumor reached 350mm$^3$. Following tumor removal, mice were continuously imaged until reaching end-point (poor body condition, PBC). (B) Tumor size of surgically removed primary tumors. (C) Bioluminescent image of metastatic burden 3 weeks post-surgery. (D) Measurement of lung photon flux (p/s) over time post-surgical removal of the primary tumor. (E) H&E quantification of spontaneous lung metastases post-surgery, taken at end-point. Data are represented as mean s.d. *t-test* was used to determine statistical significance between groups (p**<0.01). (F) Western blot depicting Gstt1 levels in *Gstt1*shRNA, i*Gstt1* cDNA or *Gstt1*shRNA + i*Gstt1* cDNA expressing primary-derived 4T1 cells with and without doxycycline. Balb/c mice were treated with doxycycline 1 week prior to orthotopic injection of *Gstt1*shRNA (n=8, -Dox), i*Gstt1* cDNA (n=4 +Dox, n=6 -Dox) or *Gstt1*shRNA + i*Gstt1* (n=7, +Dox) cDNA expressing primary-derived 4T1 cells into the mammary fat pad. Mice were monitored for primary tumor growth via bioluminescence imaging and primary tumors were surgically removed when the tumor reached 350mm$^3$. Following tumor removal, mice were continuously imaged until reaching end-point (poor body condition, PBC). (G) Weekly primary tumor growth curves prior to surgical removal using bioluminescence imaging (photon flux). (H) Representative bioluminescent image of end-point metastatic burden 4 weeks post-surgery. (I) Measurement of lung photon flux (p/s) over time post-surgical removal of the primary tumor, taken at end-point. (J) H&E quantification of spontneous lung metastases post-surgery, taken at end-point. (K) H&E quantification (Right) of spontaneous macrometastases discovered outside the lung post-surgery, taken at end-point. (L) H&E quantification of number of tissues presenting with macrometastatic tumors. Data are represented as mean s.d. *t-test* was used to determine statistical significance between groups (p**<0.01, p***<0.005, p****<0.0001).

**Supplemental Figure 4.** (A) PDAC-derived primary tumor cells expressing either Control or sh*Gstt1* were orthotopically injected into the pancreas. Mice were euthanized when primary tumor burden resulted in poor body condition (PBC), 33 days and were measured for primary tumor size. (B) Immunofluorescence staining of Gstt1 and Cytokeratin 19 (pancreatic cell marker) DTCs in orthotopic derived liver and lung tissues. (C) Immunofluorescence staining of orthotopically-derived disseminated tumor cells (DTCs) using Cytokeratin 19. (D) Quantification of single cells 33 days post-injection. Arrows indicate DTCs and inset panel demonstrates a magnified image of DTCs. Quantification represents the average of 10 fields from a minimum of n=3 mice. Data are represented as mean s.d. *t-test* was used to determine statistical significance between groups (p*<0.05).

**Supplemental Figure 5.** (A) Immunofluorescence staining of experimental disseminated tumor cells (DTCs) using Cytokeratin 19 1 week post-injection. (B) Quantification of PCNA+ and PCNA- CK19+ DTCs from (A). Quantification represents the average of 10 fields from a minimum of n=3 mice. Data are represented as mean s.d. *t-test* was used to determine statistical significance between groups (p****<0.0001). (C) FACS analysis of lung metastatic cell populations 0, 2 and 7 days post-incubation with membrane dye CM-Dil. Data represented as % of CM-Dil+ and CM-Dil- cells relative to total population.

**Supplemental Figure 6.** (A) Whole transcriptome principal component analysis (PCA) plot demonstrates distribution of mCherry$^{high}$ (n=3) and mCherry$^{low}$ (n=3) cell populations. Each sample indicates one mouse. (B) Gene set enrichment analysis (GSEA) finds hallmark enrichment of ‘Epithelial-to-Mesenchymal Transition’ and ‘TGF-Beta Signaling’ and ‘Angiogenesis’ genes in the mCherry$^{high}$ population. (C) A panel of 2D metastatic cell lines (independently derived PDAC liver and lung) expressing either Control or sh*Gstt1* were subjected to RNA-Seq (95 UP, 217 DN) (Log<2FC, FDR 0.01). DAVID biological pathway analysis on Control vs sh*Gstt1* differentially expressed gene signatures (GO_TERM_BP_DIRECT). (D) Expression (Log2CPM) of ‘Cell Cycle’ GO_TERM genes enriched in Control vs sh*Gstt1* from RNA-Seq (Log2CPM, Log<2FC, FDR 0.01). (E) Dissemination genes (Log2CPM, Hosseini et al, 2018) commonly enriched in mCherry$^{high}$ cells and DTC populations compared to mCherry$^{low}$. *t-test* was used to determine statistical significance between groups (p*<0.05). (F) Proliferation genes commonly enriched in mCherry$^{high}$ cells and DTC populations compared to mCherry$^{low}$. *t-test* was used to determine statistical significance between groups (p*<0.05).

**Supplemental Figure 7.** (A) Liver metastatic cell lines (PDAC) were lysed and subjected to immunoprecipitation using a Gstt1 antibody (whole cell lysate). Pull downs were analyzed for interactors and enriched proteins using unbiased mass spectrometry (M/S) for data in main Figure 6A. (B) Gstt1 and GSH pull downs in liver metastatic cells were blotted for validation of the interaction with Plectin. (C) Confocal imaging of immunofluorescence staining of Gstt1, Fibronectin and Cytokeratin 19 (pancreatic cell marker) in CK19$^{low}$Gstt1$^{high}$ and CK19$^{high}$Gstt1$^{low}$ liver metastatic lesions. (D) Lung metastatic cell lines (PDAC) stably expressing Control, WT iGstt1 cDNA and catalytic dead R234G iGstt1 cDNA were lysed and subjected to immunoprecipitation using a Fibronectin antibody (whole cell lysate). Pull downs were blotted for pan-glutathione (GSH), Fibronectin and Gstt1. (Right panel) Input controls. (E) mCherry$^{low}$ sorted lung metastatic cell lines stably expressing Control, WT *iGstt1* cDNA and catalytic dead R234G *iGstt1* cDNA were subjected to soft agar growth assay. (F) Quantification of soft agar colony growth in all four conditions expressed as relative to mCherry$^{low}$. The experiment was performed in triplicate sortings with three soft agar replicates each. Data are represented as mean s.e.m. *t-*

*test* was used to determine statistical significance between groups ($p^* < 0.05$). (G) Western blot depicting Gstt1 levels in WT *iGstt1* cDNA and catalytic dead R234G *iGstt1* cDNA overexpressing mCherry$^{low}$ cells. (H) Sorted mCherry$^{high}$ and mCherry$^{low}$ populations were stably transduced with sh*FN1* and subjected to soft agar growth assay. Representative images of soft agar wells. (I) Quantification of soft agar colony growth in all four conditions expressed as relative to mCherry$^{low}$. The experiment was performed in triplicate sortings with three soft agar replicates each. Data are represented as mean s.e.m. *t-test* was used to determine statistical significance between groups ($p^* < 0.05$; $p^{**} < 0.01$). (J) Western blot analysis of conditions in (H) and (I).

### Supp Fig 1.

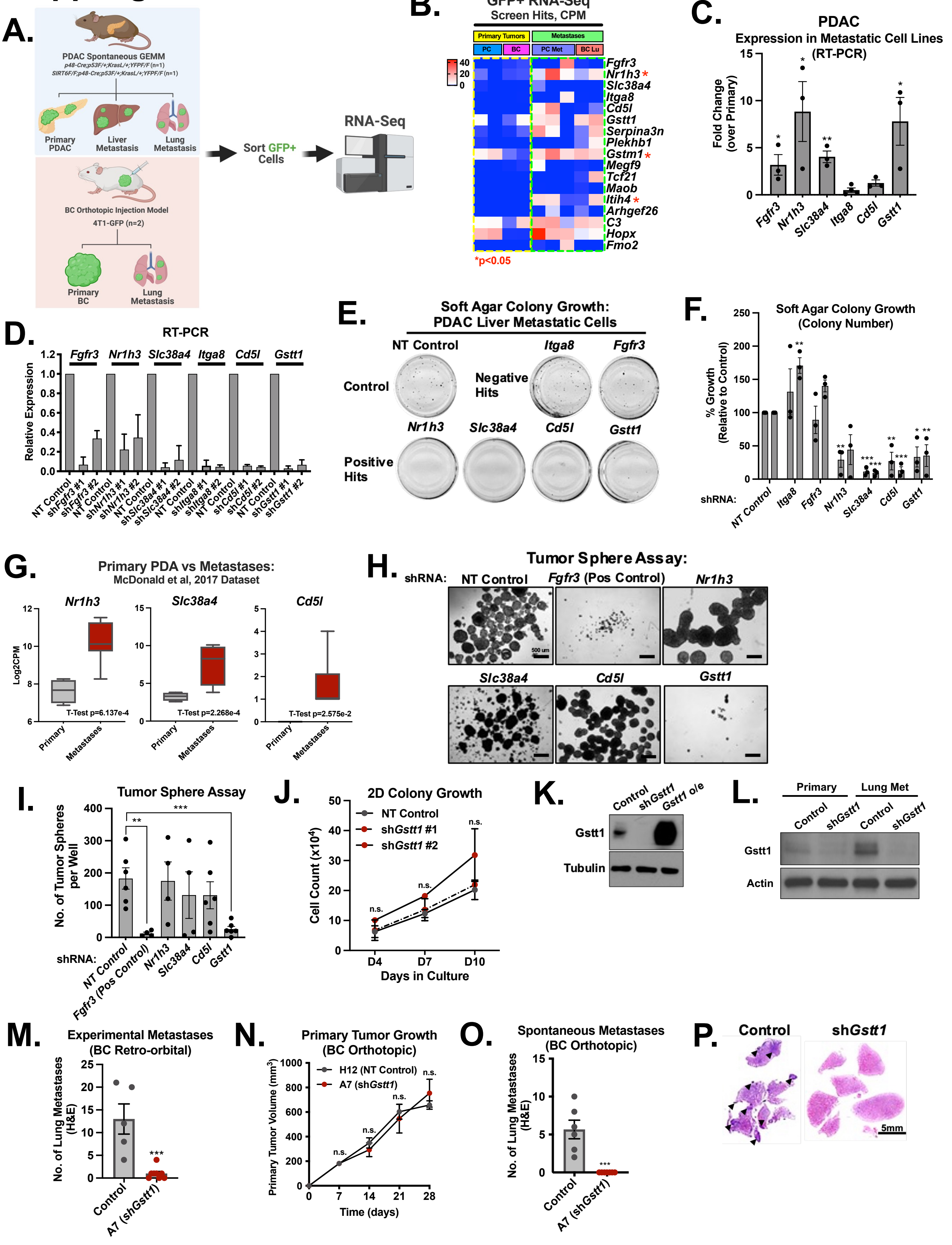

A.
PDAC Spontaneous GEMM
Primary PDAC
Liver Metastasis
Lung Metastasis
BC Orthotopic Injection Model
4T1-GFP (n=2)
Primary BC
Lung Metastasis
Sort GFP+ Cells
RNA-Seq
B.
GFP+ RNA-Seq
Screen Hits, CPM
Primary Tumors
Metastases
PC
BC
PC Met
BC Lu
Fgfr3
Nr1h3 *
Slc38a4
Itga8
Cd5l
Gstt1
Serpina3n
Plekhb1
Gstm1 *
Megf9
Tcf21
Maob
Itih4 *
Arhgef26
C3
Hopx
Fmo2
*p<0.05
C.
PDAC
Expression in Metastatic Cell Lines (RT-PCR)
Fold Change (over Primary)
D.
RT-PCR
Relative Expression
E.
Soft Agar Colony Growth: PDAC Liver Metastatic Cells
NT Control
Control
Negative Hits
Positive Hits
F.
Soft Agar Colony Growth (Colony Number)
% Growth (Relative to Control)
shRNA:
G.
Primary PDA vs Metastases:
McDonald et al, 2017 Dataset
Log2CPM
T-Test p=6.137e-4
T-Test p=2.268e-4
T-Test p=2.575e-2
Primary
Metastases
H.
Tumor Sphere Assay:
shRNA:
Fgfr3 (Pos Control)
I.
Tumor Sphere Assay
No. of Tumor Spheres per Well
Fgfr3 (Pos Control)
J.
2D Colony Growth
NT Control
shGstt1 #1
shGstt1 #2
Cell Count (x10⁴)
Days in Culture
K.
Control
shGstt1
Gstt1 o/e
Gstt1
Tubulin
L.
Primary
Lung Met
Actin
M.
Experimental Metastases (BC Retro-orbital)
No. of Lung Metastases (H&E)
A7 (shGstt1)
N.
Primary Tumor Growth (BC Orthotopic)
H12 (NT Control)
A7 (shGstt1)
Primary Tumor Volume (mm³)
Time (days)
O.
Spontaneous Metastases (BC Orthotopic)
P.
5mm

### Supp Fig 2.

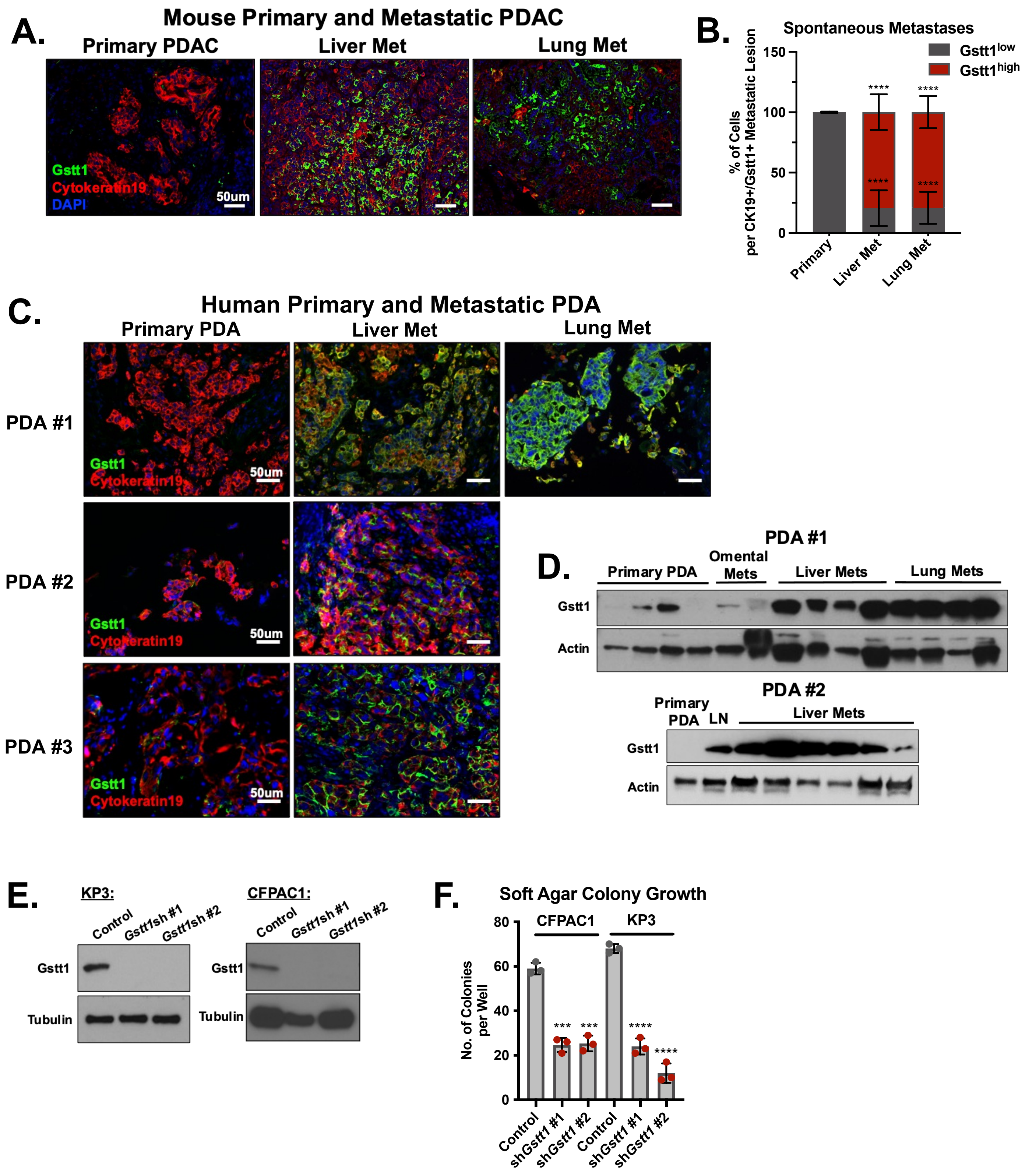

A.
Mouse Primary and Metastatic PDAC
Primary PDAC
Liver Met
Lung Met
Gstt1
Cytokeratin19
DAPI
50um
B.
Spontaneous Metastases
Gstt1low
Gstt1high
% of Cells per CK19+/Gstt1+ Metastatic Lesion
150
100
50
0
****
Primary
Liver Met
Lung Met
C.
Human Primary and Metastatic PDA
Primary PDA
Liver Met
Lung Met
PDA #1
PDA #2
PDA #3
Gstt1
Cytokeratin19
50um
D.
PDA #1
Primary PDA
Omental Mets
Liver Mets
Lung Mets
Gstt1
Actin
PDA #2
Primary PDA
LN
Liver Mets
Gstt1
Actin
E.
KP3:
CFPAC1:
Control
Gstt1sh #1
Gstt1sh #2
Gstt1
Tubulin
F.
Soft Agar Colony Growth
CFPAC1
KP3
No. of Colonies per Well
80
60
40
20
0
***
****
Control
shGstt1 #1
shGstt1 #2

### Supp Fig 3.

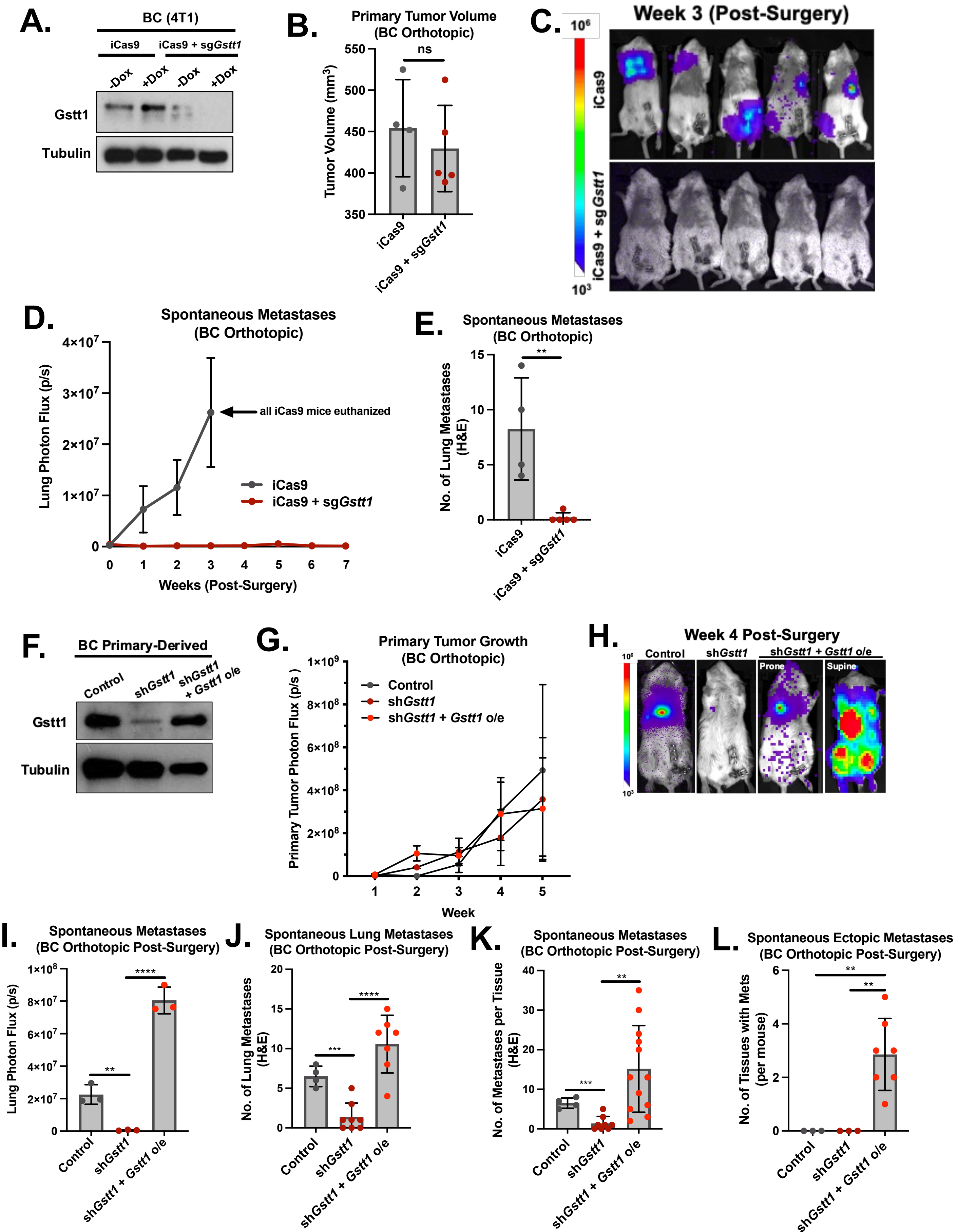

A.
BC (4T1)
iCas9
iCas9 + sgGstt1
-Dox
+Dox
-Dox
+Dox
Gstt1
Tubulin
B.
Primary Tumor Volume
(BC Orthotopic)
ns
Tumor Volume (mm³)
550
500
450
400
350
iCas9
iCas9 + sgGstt1
C.
Week 3 (Post-Surgery)
10⁶
10³
iCas9
iCas9 + sg Gstt1
D.
Spontaneous Metastases
(BC Orthotopic)
Lung Photon Flux (p/s)
4×10⁷
3×10⁷
2×10⁷
1×10⁷
0
all iCas9 mice euthanized
iCas9
iCas9 + sgGstt1
0 1 2 3 4 5 6 7
Weeks (Post-Surgery)
E.
Spontaneous Metastases
(BC Orthotopic)
No. of Lung Metastases
(H&E)
15
10
5
0
**
iCas9
iCas9 + sgGstt1
F.
BC Primary-Derived
Control
shGstt1
shGstt1
+ Gstt1 o/e
Gstt1
Tubulin
G.
Primary Tumor Growth
(BC Orthotopic)
Primary Tumor Photon Flux (p/s)
1×10⁹
8×10⁸
6×10⁸
4×10⁸
2×10⁸
0
Control
shGstt1
shGstt1 + Gstt1 o/e
1 2 3 4 5
Week
H.
Week 4 Post-Surgery
10⁶
10³
Control
shGstt1
shGstt1 + Gstt1 o/e
Prone
Supine
I.
Spontaneous Metastases
(BC Orthotopic Post-Surgery)
Lung Photon Flux (p/s)
1×10⁸
8×10⁷
6×10⁷
4×10⁷
2×10⁷
0
**
****
Control
shGstt1
shGstt1 + Gstt1 o/e
J.
Spontaneous Lung Metastases
(BC Orthotopic Post-Surgery)
No. of Lung Metastases
(H&E)
20
15
10
5
0
***
****
Control
shGstt1
shGstt1 + Gstt1 o/e
K.
Spontaneous Metastases
(BC Orthotopic Post-Surgery)
No. of Metastases per Tissue
(H&E)
40
30
20
10
0
***
**
Control
shGstt1
shGstt1 + Gstt1 o/e
L.
Spontaneous Ectopic Metastases
(BC Orthotopic Post-Surgery)
No. of Tissues with Mets
(per mouse)
6
4
2
0
**
**
Control
shGstt1
shGstt1 + Gstt1 o/e

### Supp Fig 4.

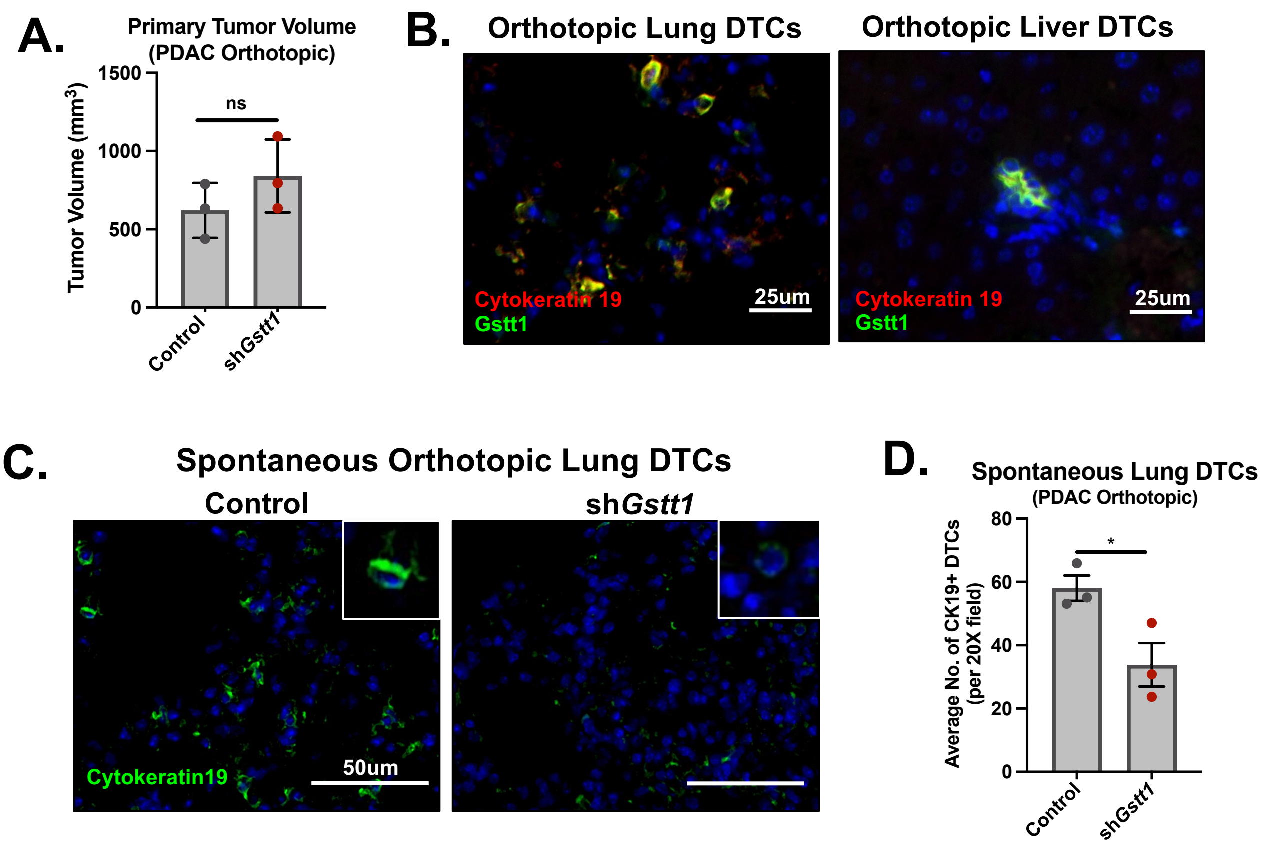

A.
Primary Tumor Volume
(PDAC Orthotopic)
Tumor Volume (mm³)
1500
1000
500
0
ns
Control
shGstt1
B.
Orthotopic Lung DTCs
Orthotopic Liver DTCs
Cytokeratin 19
Gstt1
25um
C.
Spontaneous Orthotopic Lung DTCs
Control
shGstt1
Cytokeratin19
50um
D.
Spontaneous Lung DTCs
(PDAC Orthotopic)
Average No. of CK19+ DTCs
(per 20X field)
80
60
40
20
0
*

### Supp Fig 5.

A.

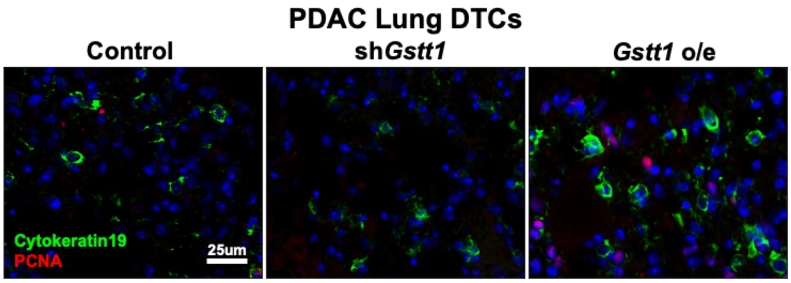


B.

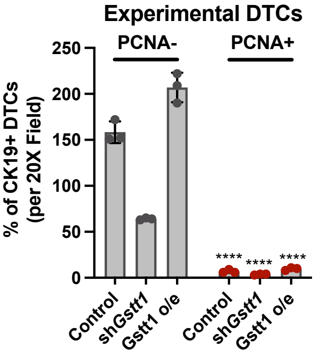


C.

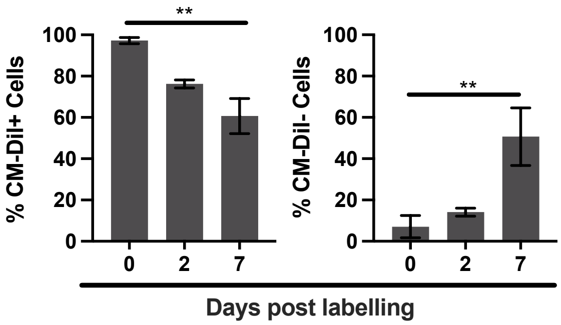

### Supp Fig 6.

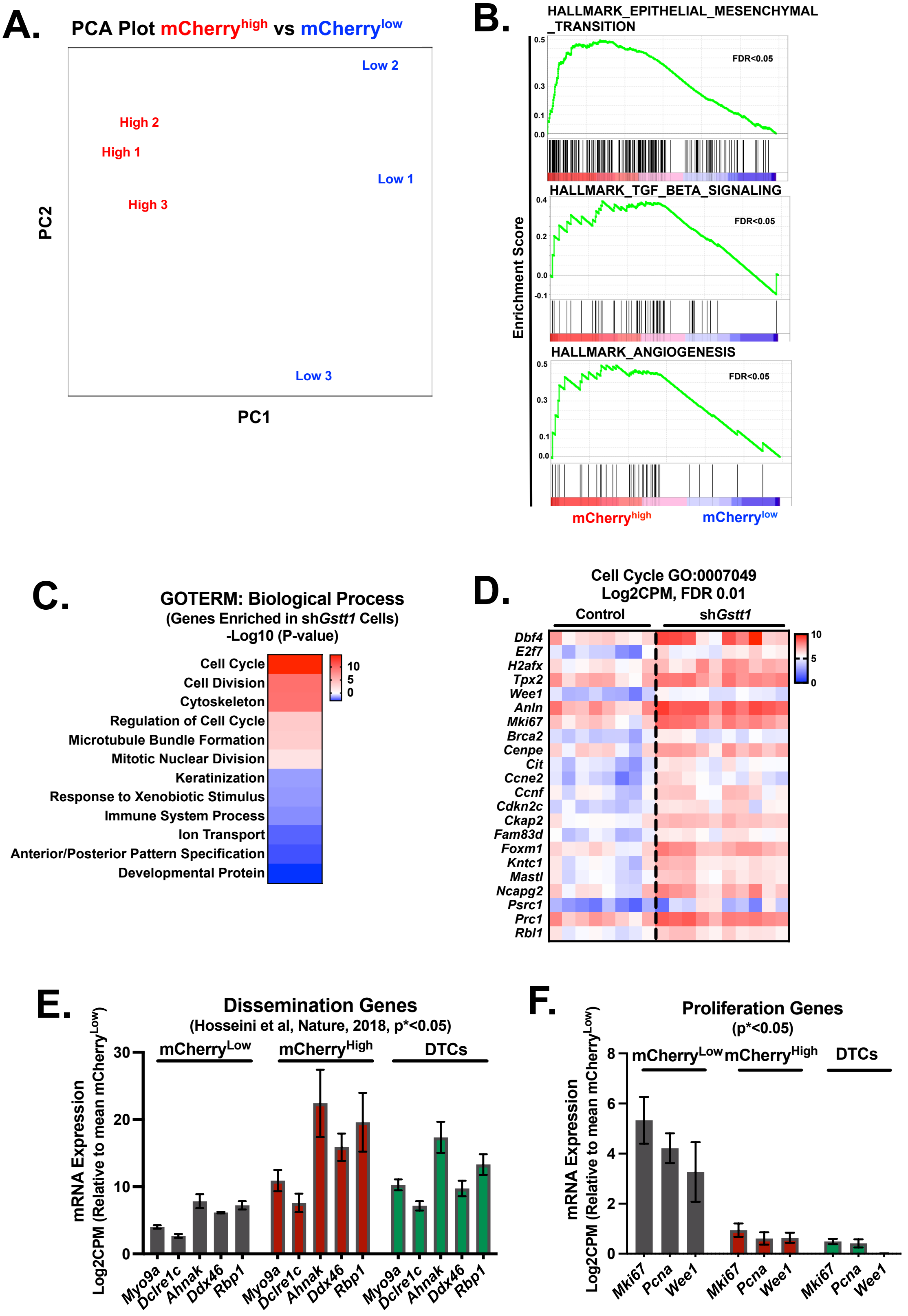
A.
PCA Plot mCherry^high vs mCherry^low
Low 2
High 2
High 1
Low 1
High 3
Low 3
PC2
PC1
B.
HALLMARK_EPITHELIAL_MESENCHYMAL_TRANSITION
FDR<0.05
HALLMARK_TGF_BETA_SIGNALING
FDR<0.05
HALLMARK_ANGIOGENESIS
FDR<0.05
Enrichment Score
mCherry^high
mCherry^low
C.
GOTERM: Biological Process
(Genes Enriched in shGstt1 Cells)
-Log10 (P-value)
Cell Cycle
Cell Division
Cytoskeleton
Regulation of Cell Cycle
Microtubule Bundle Formation
Mitotic Nuclear Division
Keratinization
Response to Xenobiotic Stimulus
Immune System Process
Ion Transport
Anterior/Posterior Pattern Specification
Developmental Protein
D.
Cell Cycle GO:0007049
Log2CPM, FDR 0.01
Control
shGstt1
Dbf4
E2f7
H2afx
Tpx2
Wee1
Anln
Mki67
Brca2
Cenpe
Cit
Ccne2
Ccnf
Cdkn2c
Ckap2
Fam83d
Foxm1
Kntc1
Mastl
Ncapg2
Psrc1
Prc1
Rbl1
E.
Dissemination Genes
(Hosseini et al, Nature, 2018, p*<0.05)
mCherry^Low
mCherry^High
DTCs
mRNA Expression
Log2CPM (Relative to mean mCherry^Low)
Myo9a
Dclre1c
Ahnak
Ddx46
Rbp1
F.
Proliferation Genes
(p*<0.05)
mCherry^Low
mCherry^High
DTCs
Mki67
Pcna
Wee1

### Supp Fig 7.

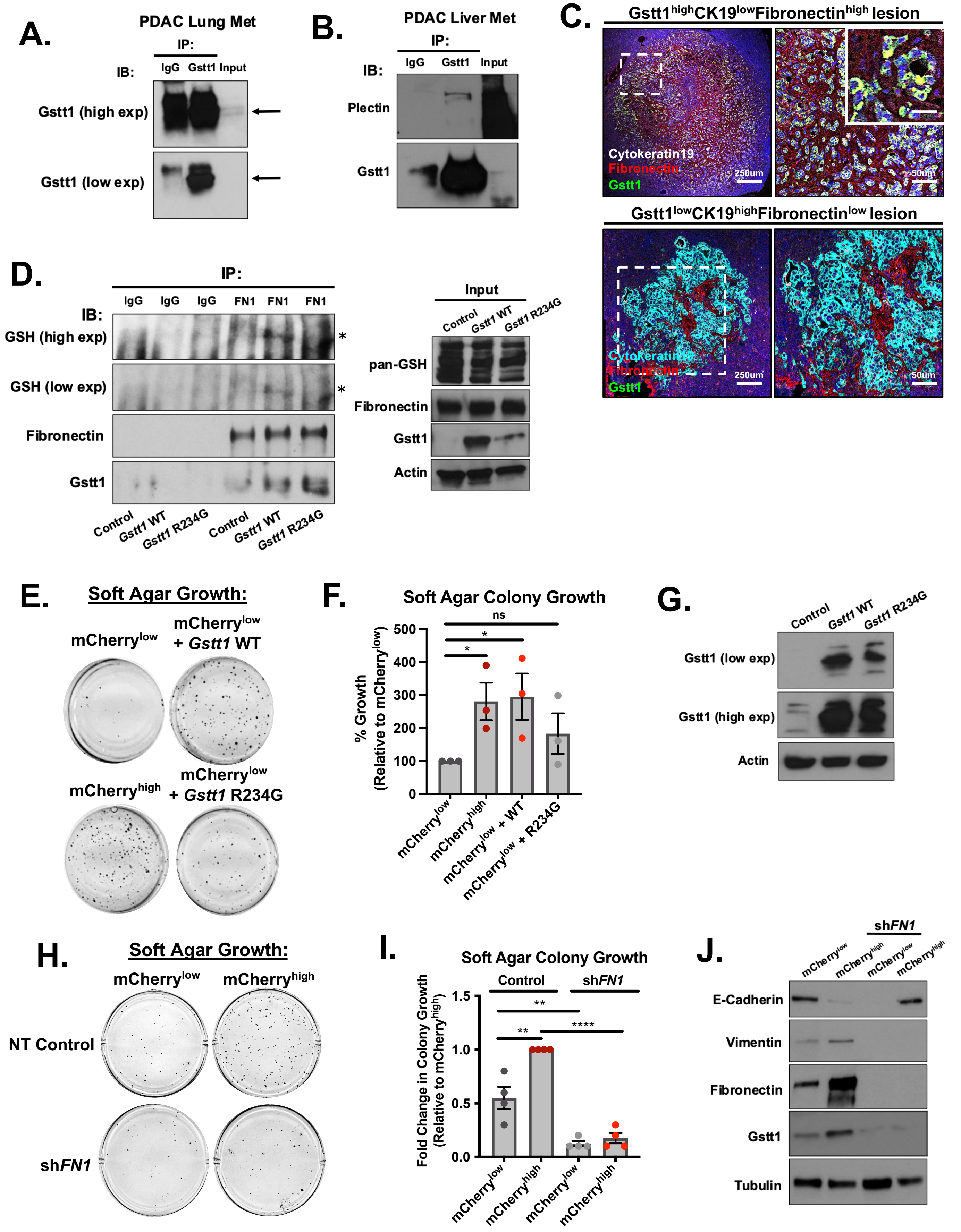

A.
PDAC Lung Met
IP:
IB:
IgG
Gstt1
Input
Gstt1 (high exp)
Gstt1 (low exp)
B.
PDAC Liver Met
IP:
IB:
IgG
Gstt1
Input
Plectin
Gstt1
C.
Gstt1highCK19lowFibronectinhigh lesion
Cytokeratin19
Fibronectin
Gstt1
250um
50um
Gstt1lowCK19highFibronectinlow lesion
Cytokeratin19
Fibronectin
Gstt1
250um
50um
D.
IP:
IB:
IgG
IgG
IgG
FN1
FN1
FN1
GSH (high exp)
GSH (low exp)
Fibronectin
Gstt1
Control
Gstt1 WT
Gstt1 R234G
Control
Gstt1 WT
Gstt1 R234G
*
*
Input
Control
Gstt1 WT
Gstt1 R234G
pan-GSH
Fibronectin
Gstt1
Actin
E.
Soft Agar Growth:
mCherrylow
mCherrylow + Gstt1 WT
mCherryhigh
mCherrylow + Gstt1 R234G
F.
Soft Agar Colony Growth
ns
*
*
% Growth (Relative to mCherrylow)
500
400
300
200
100
0
mCherrylow
mCherryhigh
mCherrylow + WT
mCherrylow + R234G
G.
Control
Gstt1 WT
Gstt1 R234G
Gstt1 (low exp)
Gstt1 (high exp)
Actin
H.
Soft Agar Growth:
mCherrylow
mCherryhigh
NT Control
shFN1
I.
Soft Agar Colony Growth
Control
shFN1
**
**
****
Fold Change in Colony Growth (Relative to mCherryhigh)
1.5
1.0
0.5
0.0
mCherrylow
mCherryhigh
mCherrylow
mCherryhigh
J.
shFN1
mCherrylow
mCherryhigh
mCherrylow
mCherryhigh
E-Cadherin
Vimentin
Fibronectin
Gstt1
Tubulin
